# The 3D genome organization of *Drosophila melanogaster* through data integration

**DOI:** 10.1101/099911

**Authors:** Qingjiao Li, Harianto Tjong, Xiao Li, Ke Gong, Xianghong Jasmine Zhou, Irene Chiolo, Frank Alber

**Author notes:** Correspondence should be addressed to F.A. and I.C. These authors contributed equally (Qingjiao Li, Harianto Tjong).

## Abstract

Genome structures are dynamic and non-randomly organized in the nucleus of higher eukaryotes. To maximize the accuracy and coverage of 3D genome structural models, it is important to integrate all available sources of experimental information about a genome’s organization. It remains a major challenge to integrate such data from various complementary experimental methods. Here, we present an approach for data integration to determine a population of complete 3D genome structures that are statistically consistent with data from both genome-wide chromosome conformation capture (Hi-C) and lamina-DamID experiments. Our structures resolve the genome at the resolution of topological domains, and reproduce simultaneously both sets of experimental data. Importantly, this framework allows for structural heterogeneity between cells, and hence accounts for the expected plasticity of genome structures. As a case study we choose *Drosophila melanogaster* embryonic cells, for which both data types are available. Our 3D geome structures have strong predictive power for structural features not directly visible in the initial data sets, and reproduce experimental hallmarks of the *D. melanogaster* genome organization from independent and our own imaging experiments. Also they reveal a number of new insights about the genome organization and its functional relevance, including the preferred locations of heterochromatic satellites of differnet chromosomes, and observations about homologous pairing that cannot be directly observed in the original Hi-C or lamina-DamID data. To our knowledge our approach is the first that allows systematic integration of Hi-C and lamina DamID data for complete 3D genome structure calculation, while also explicitly considering genome structural variability.

## Introduction

It has become increasingly clear that a chromosome’s three-dimensional organization influences the regulation of gene expression and other genome functions. Early microscopy and biochemical studies showed that chromosomes in higher eukaryotes form distinct territories, which although stochastically organized tend to be located at preferred positions within the nucleus. For example, lamina-DamID experiments have identified specific chromatin domains with a high propensity to be located at the nuclear envelope (NE), confirming the important role of the NE in spatial genome organization and gene regulation in Drosophila, human and mouse [1–3]. Chromosome conformation capture experiments (Hi-C and variants) detect chromatin interactions at genome-wide scale [4–10] and reveal a hierarchical chromosome organization: the chromatin can be segmented into domains, which in turn combine to form subcompartments of functionally related chromatin [10–12]. Topological associated domains (TADs) are defined by observing an increased probability of interaction between chromatin regions in a domain relative to interactions between domains. In addition, it has been shown that the border regions between domains are enriched in specific insulator proteins, such as CTCF and ZNF143 in mammalian cells and BEAF, CTCF and CP190 in Drosophila cells. However, the precision of domain border detection depend to some extent on the sequencing depth as well as algorithmic parameter settings. At increased sequencing depth it is possible to detect reliably individual chromatin loops, which often demarcate contact domains (at ~100kb domain length) [5].

Computational approaches can aid in mapping the global 3D structures of genomes at various scales [13–19]. These are currently divided into data-driven and *de novo* modeling techniques [20]. Data-driven models use experimental information, often Hi-C data, to generate 3D genome structures that are constrained to be consistent with the data. Data-driven models can be further subdivided into two classes. The first represents the genome as a consensus structure most consistent with the data. This approach often assumes an anti-correlation between the Hi-C contact probability of two chromatin regions and their average spatial distance. In contrast, the second class of methdos explicitly models the large variability of genome structures between isogenic cells (even within a sample of synchronized cells) by creating a population of thousands of model structures, in which the accumulated chromatin contacts in all structures reproduce the observed Hi-C matrix rather than each structure individually. These approaches do not need to assume any functional relationship between contact frequencies and spatial distances.

We introduced a method for population-based modeling to analyze the structure of complete diploid genomes from Hi-C data [6, 21, 22]. Our method uses an iterative, probabilistic optimization framework to deconvolve the Hi-C data into a population of individual structures by inferring cooperative chromatin interactions that are likely to co-occur in the same cells. Our method generates a large number of genome structures whose chromatin contacts in the models are statistically consistent with those from the Hi-C data. Other ensemble-based methods have been introduced and applied to individual chromosomes or chromatin domains [13, 19].

So far, computational models of genome structures have typically relied on just one data type, such as Hi-C, even though a single experimental method cannot capture all aspects of the spatial genome organization. However, data are available from a wide range of technologies with complementary strengths and limitations. Integrating all these different data types would greatly increase the accuracy and coverage of genome structure models. Moreover, such models would offer a way to cross-validate the consistency of data obtained from complementary technologies. For example, lamina-DamID experiments show a chromatin region’s probability to be close to the lamina at the nuclear envelope [23, 24], while Hi-C experiments reveal the probability that two chromatin regions are in spatial proximity. Large-scale 3D fluorescence in situ hybridization (FISH) experiments show the distance between loci directly, and can be used to measure the distribution of distances across a population of cells.

It remains a major challenge to develop hybrid methods that can systematically integrate data from many different technologies to generate structural maps of the genome. In this paper, we present a method for integrating data from Hi-C and lamina-DamID experiments to maximize the accuracy of population-based 3D genome structural models. We apply this approach to model the diploid genome of Drosophila.

*Drosophila melanogaster* is a popular model organism to study the organization and functional relevance of 3D genome structure, owing to its relative small genome and the availability of many genetic tools. A variety of microscopy-based experiments have already studied the nuclear organization of *D. melanogaster*, and elucidated some regulatory mechanisms [25–29]. For example, the pairing of homologous chromosomes has been observed in the somatic cells of *D. melanogaster* and other dipteran insects [30–33]. This kind of pairing can influence gene expression by forming interactions between regulatory elements on homologous chromosomes, a process called transvection [26, 34]. Although transvection is common in Drosophila, not every gene region with homologue pairing shows transvection. Therefore, questions remain as to whether somatic homolog pairing has other regulatory roles. In Drosophila, the centromeres tend to cluster and relocate close to the periphery of the nucleolus during interphase [35]. Centromere clustering is also observed in many other organisms, including yeast, mouse and human, and this process is thought to play an important role in determining the overall genome architecture [36, 37].

Over the past ten years, high-throughput genetic and genomic techniques have generated genome-wide maps of histone modifications, transcription factor binding, and chromatin interactions for *D. melanogaster* [1, 7, 8, 38, 39]. Pickersgill *et al.* used lamina-DamID experiments combined with a microarray technique to detect the binding signals of genome-wide chromatin to the lamina matrix [1]. Around 500 genes were detected to interact with the lamina. These genes were transcriptionally silenced and late-replicating. Pickersgill *et al.* then used FISH experiments to confirm that the lamina-targeted loci were more frequently located at the nuclear envelope than other loci. Recently, genome-wide chromatin contacts have been determined for 16-18 hr Drosophila embryos using the Hi-C technique [8]. The euchromatin genome was divided into 1169 physical domains based on Hi-C interaction profiles. These physical domains (which would be referred to as topological associated domain, or TADs, in mammalian cells) were assigned to four functional classes based on their average epigenetic signatures: Null, Active, Polycomb-Group (Pc-G) and HP1/Centromere.

Despite all this work, the global 3D nuclear architecture of the *D. melanogaster* genome is still unknown. Because both Hi-C and lamina-DamID data are available for Drosophila embryonic cells, we use these data to test our integration method. Each diploid genome structure in our population-based model is defined by the 3D positions of all 1169 TADs. The structures are generated by optimizing a likelihood function, so that the ensemble is statistically consistent with both the experimentally derived contact probabilities between all chromatin domains from Hi-C data and the probability that a given chromatin domain is close to the NE from lamina-DamID data.

We validated our 3D genome models against independent experimental data and known structural features. Our models confirm the formation of distinct chromosome territories, with relatively low rates of intermingling between chromosomes [40, 41]. In addition, our models often show a polarized organization of chromosomes in the nucleus [27, 42, 43]. Analysis of the model population leads to a number of new insights about the nuclear organization of *D. melanogaster* and its functional relevance. For instance, our models reveal the preferred locations of heterochromatin and the nucleolus, which we were able to confirm by 3D FISH experiments. The nucleolus serves as an anchor for chromosomes, and is surrounded by pericentromeric heterochromatin. The distance of pericentromeric heterochromatin regions from the periphery varies by chromosome, with chromosomes 4 and X heterochomatin more peripheral relative to Pericentromeric regions of other chromosomes. Interestingly, the frequency of homologous pairing varies along the chromosomes with the lowest frequencies observed in our models for domains enriched in protein binding sites for Mrg15. These observations support the model that Mrg15 plays a role in the dissociation of homologous chromosome pairs during interphase, as previously suggested [44]. Finally, the structure population suggests that homologous chromosome pairing plays a functional role in transcriptional activity and DNA replication program.

## Results

### Population-based genome structure modeling from data integration

Our goal is to determine a population of 3D genome structures for *Drosophila melanogaster* that is consistent with data from Hi-C and lamina-DamID experiments. Suppose A is a probability matrix derived from Hi-C data, and *E* is a probability vector derived from lamina-DamID data. The elements of A describe how frequently a given pair of TADs are in contact with each other in an ensemble of cells, and *E* describes how frequently a given TAD is in contact with the nuclear envelope (NE). The goal is to generate a population of genome structures **X**, whose TAD-TAD and TAD-NE contact frequencies are statistically consistent with both **A** and *E*. We formulate the genome structure modeling problem as a maximization of the likelihood *P*(**A**,*E*|**X**).

More specifically, the structure population is defined as a set of *M* diploid genome structures **X** = {**X**_1_,**X**_2_,…,**X**_*M*_}, where the *m*-th structure **X**_*m*_ is a set of 3-dimensional vectors representing the center coordinates of 2*N* domain spheres 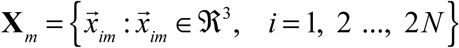. *N* is the number of TAD domains, but each domain has two homologous copies with different coordinates. The contact probability matrix **A** = (*a_IJ_*)_*N×N*_ for *N* domains is derived from the Hi-C data, which do not distinguish between homologous copies. Each element *a_IJ_* is the probability that a direct contact between domains *I* and *J* exists in a structure of the population. (Note that the capital letter indices *I* and *J* refer to domains without distinguishing between their homologous copies, while the lowercase indices *i*, *i’* and *j*, *j’* do distinguish between copies.) The contact probability vector *E* = {*e_I_*|*I* = 1,2,…,*N*} is derived from the lamina-DamID data, and defines the probability for each TAD to be localized at the nuclear envelope (NE). With known **A** and *E*, we calculate the structure population **X** such that the likelihood *P* (**A**,*E*|**X**) is maximized.

The Hi-C and lamina-DamID experiments provide data that is averaged over a large population of cells, so they cannot reveal which contacts co-exist in the same 3D structure. Therefore, both **A** and *E* are interpreted as ensemble averages. To represent information derived from individual cells, we introduce two latent variables **W** and **V**. The “contact indicator tensor” **W** = (*w_ijm_*)_2*N*×2*N*×*M*_ is a binary, 3rd-order tensor. It contains the information missing from the Hi-C data **A**, namely which domain contacts belong to each of the *M* structures in the model population and also which homologous chromosome copies are involved ( *w_ijm_* = 1 indicates a contact between domain spheres *i* and *j* in structure m; *w_ijm_* = 0 otherwise). **W** is a detailed expansion of **A** into a diploid, single-structure representation of the data. The structure population **X** is consistent with **W**. Therefore, the dependence relationship between these three variables is given as **X** → **W** → **A**. Another latent variable, **V** = (*v_im_*)_2*N*×*M*_, specifies which domain is located near the NE in each structure of the population and also distinguishes between the two homologous TAD copies (*v_im_* = 1 indicates that TAD *i* is located near the NE in structure *m*; *v_im_* = 0 otherwise). The dependence relationship between **X**, **V** and *E* is given as **X** → **V** → *E*, because **X** is the structure population consistent with **V** and **V** is a detailed expansion of *E* at a diploid and single-structure representation of the data.

In addition to the Hi-C and lamina-DamID data, we also consider additional information specific for Drosophila genome organization, e.g. the nuclear volume, an upper bound for homolog chromosome pairing, constraints connecting consecutive domains (including heterochromatin domains) as well as costraints for anchoring centromeres to the nucleolus (see the detailed description in the **Materials and Methods** section).

Thus, the optimization problem is expressed as:

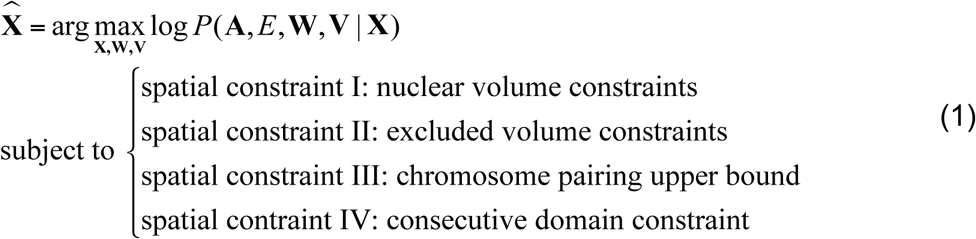

The log likelihood can be expanded as

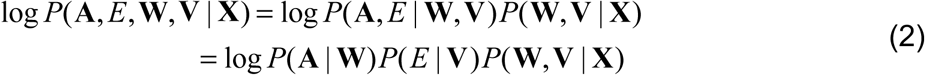

We have developed a variant of the EM method to iteratively optimize the log likelihood [22]. Each iteration consists of two steps (**Fig. 1A**):

**Figure 1.**
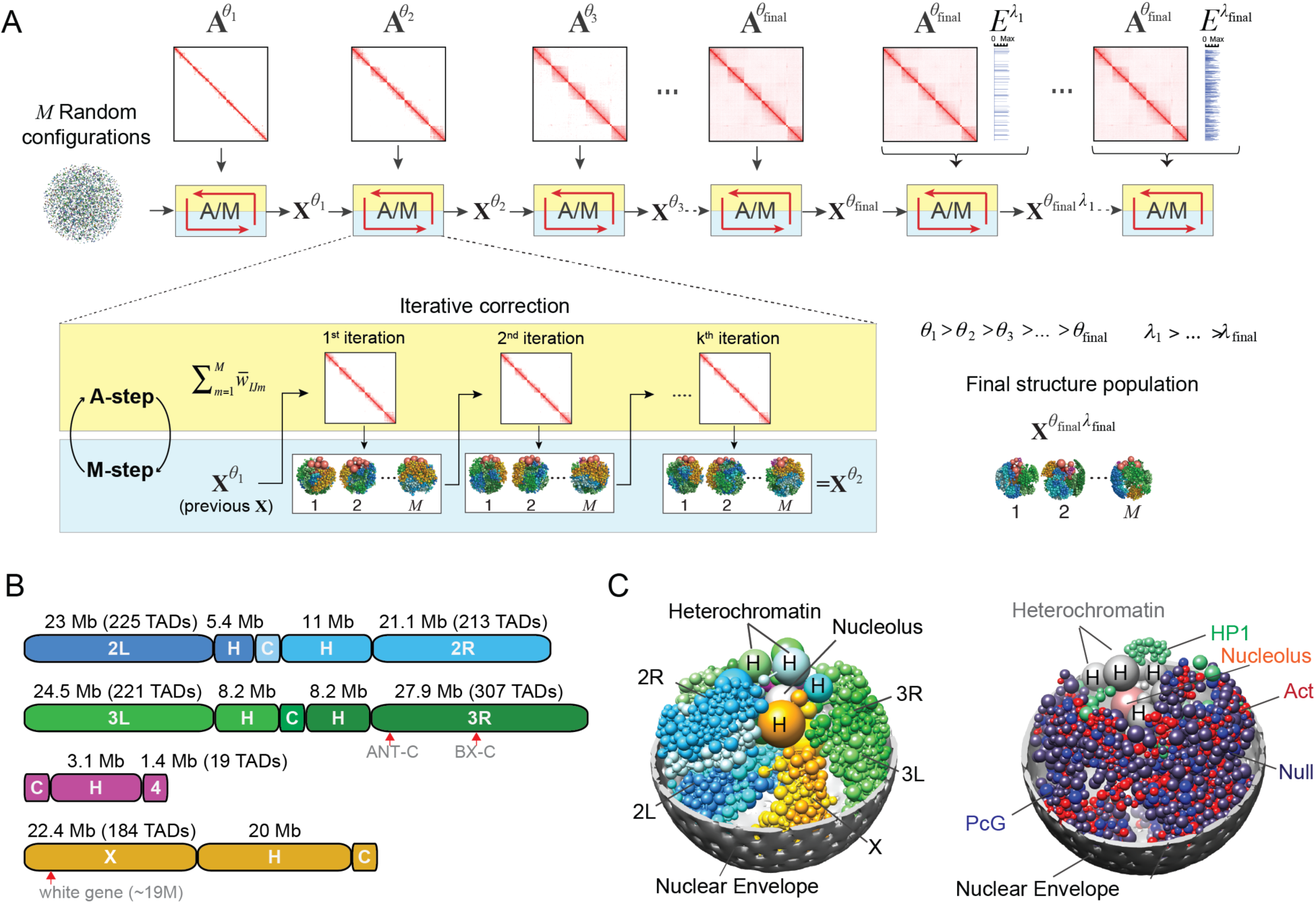
Overview of the population-based genome structure modeling approach and its application to the Drosophila genome. (A) The initial structures are random configurations. Maximum likelihood optimization is achieved through an iterative process with two steps, assignment (A) and modeling (M). We increase the optimization hardness over several stages by including contacts from the Hi-C matrix **A** with lower probability thresholds (*θ*). After the population reproduces the complete Hi-C data, we include the vector *E* (lamina-DamID), again in stages with decreasing contact probability thresholds (*λ*). (B) Schematic of the Drosophila genome. The autosome arms are designated 2L, 2R, 3L, 3R, 4 and X. The arms of chr2 and chr3 are connected by centromeres labeled “C”. Euchromatic regions are labeled as the arm. The numbers along the top of a genome indicate the length of the section in megabases (Mb), and for euchromatin the number of spheres (TADs) in the structure model is also given. The heterochromatic region of each chromosome arm is labeled “H”. The white gene is located ~19M away from the heterochromatin of chrX. Also indicated are the Hox genes: 5 genes of the Antennapedia complex (ANT-C) are located at ~2.3M-2.8M from the heterochromatin of chr3R, and 3 genes of the Bithorax complex (BX-C) are located at ~12.4M-12.7M from the heterochromatin of chr3R. (C) Snapshot of a single structure randomly picked from the final population. (Left panel) The full diploid chromosomes are shown in colors: blue-chr2, green-chr3, magenta-chr4, and orange-chrX. The heterochromatin spheres are larger than the euchromatin domains. The nucleolus is colored in silver. (Right panel) The euchromatin domains are colored to reflect their epigenetic class: red-Active, blue-PcG, green-HP1 and dark-Null. Heterochromatin spheres are grey, and the nucleolus is pink.

- Assignment step (*A-step*): Given the current model **X**^(*i*)^, estimate the latent variables **W**^(*i*+1)^ and **V**^(*i*+1)^ by maximizing the log-likelihood over all possible values of **W** and **V**.

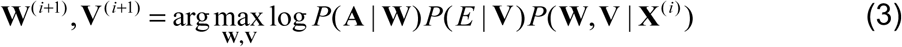
- Modeling step (*M-step*): Given the current estimated latent variables **W**^(*i*+1)^ and **V**^(*i*+1)^, find the model **X**^(*i*+1)^ that maximizes the log-likelihood function.

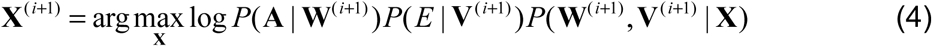 The detailed implementation of the A-step and M-step are described in **Materials and Methods**. We follow the step-wise optimization strategy described previously [22], and gradually increase the optimization hardness by adding contact constraints at a decreasing contact probability threshold.

### A population of Drosophila genome structures at the TAD level

The euchromatin regions of *Drosophila melanogaster* chromosomes 2, 3, 4, and X are partitioned into 1169 TADs, as previously described [8]. The region of pericentromeric heterochromatin of each chromosome arm is spatially clustered and represented by a single domain (**Fig. 1B**) [45–47] (**Materials and Methods**). The nuclear diameter is set to 4 microns. The model also contains a nucleolus, represented by a sphere with a radius 1/6 of the nuclear radius. We estimated the nucleolus volume from our immunofluorescence analysis of Drosophila Kc cells (**Fig. S7A**) (**Materials and Methods**).

By optimizing the likelihood function (Eq. (1)) we generated a population of 10,000 genome structures that accurately reproduces the domain contact probabilities from Hi-C experiments and the probabilities for domains to reside at the nuclear envelope (NE) from lamina-DamID experiments (**Materials and Methods**). For comparison, we also generated a population of structures using only Hi-C data, referred to hereafter as a control model. To test the reproducibility of our method, we generated a second, independently calculated model. The second model confirms our conclusions (**Figs. 6C and S10**).

### Validation of the structure population

#### Reproducing the Hi-C contact probabilities

We first assessed the consistency between the chromatin contact probabilities in our structure population with those observed experimentally. The contact probability of any two domains is defined as the fraction of model genome structures for which the two domains are in physical contact with each other,measured over the entire population (a domain-domain contact is defined by an overlap between their soft sphere contact radius). The domain contact probability matrix in our model shows excellent agreement (high correlation) with the Hi-C data, and also closely reproduces the interaction patterns visible in the matrix. The average column-based Pearson’s correlation coefficient (PCC) is 0.9840, and the element-wise PCC is 0.9839 (**Suppl. Table 1**). The correlation coefficients of the intra-chromosome arm contact probabilities range between 0.9795 and 0.9981 over all arms, confirming the excellent visual comparison shown in **Fig. 2A**. The correlation coefficients for inter-arm and inter-chromosome contact probabilities are lower, ranging between 0.1475 and 0.3822 (**Suppl. Table 1**). This relatively weak agreement between the model and the experimental data for inter-arm and inter-chromosome interactions can be explained by the following argument. In the Hi-C data, inter-arm and inter-chromosome interactions are relatively infrequent and unstructured, indicating that contacts between chromosomes are predominantly random. Due to their low occurrence, these interactions are also less reproducible than intra-arm interactions, especially at low sequencing depth. This reasoning is confirmed by comparing two Hi-C experiments performed with two different restriction enzymes [6, 48]. The differences in contact frequencies between the two experiments are generally much larger for interchromosome arm interactions than for intra-chromosome arm interactions.

**Figure 2.**
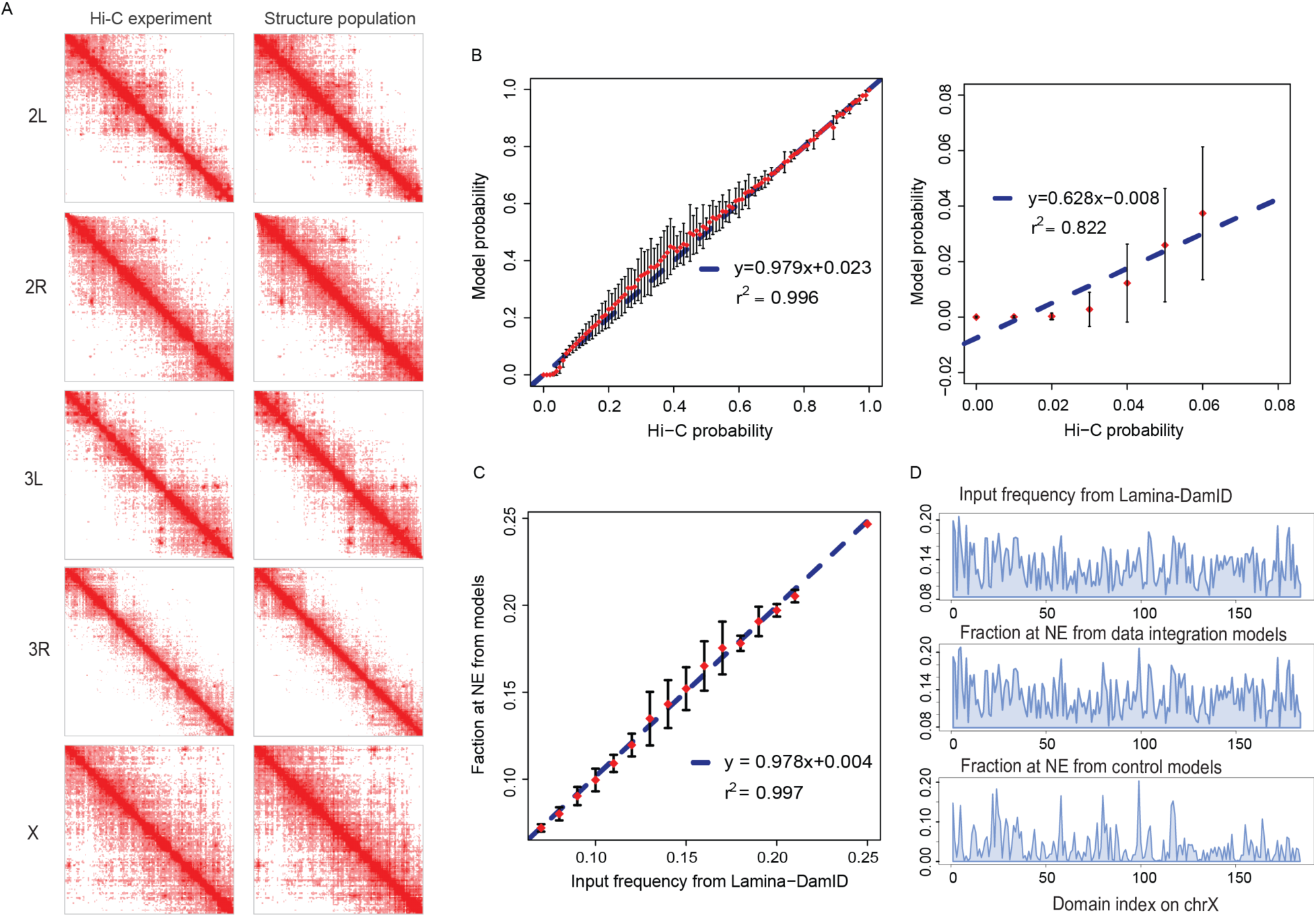
Reproduction of Hi-C and lamina-DamID data. (A) Heat maps of intra-arm contact probabilities from Hi-C experiments (left) and intra-arm contact frequencies from the structure population (right). Their similarity is quantified by element-wise Pearson’s correlations, which are 0.9844, 0.9852, 0.9840, 0.9859 and 0.9795 for chr2L, chr2R, chr3L, chr3R and chrX, respectively. The maps only show interactions with probabilities no less than 6%, which are used as constraints in our modeling procedure. (B) Agreement between the experimental data and model contact probabilities. (Left panel) The input Hi-C probabilities are divided into 100 bins, the corresponding model probabilities in one bin are summarized by mean and variance, and then the error bar plot is shown. The blue dot-line is the linear regression line between the average model probabilities of each bin and the mid-point Hi-C probabilities of the bins. Their Pearson’s correlation is 0.998 with p-value < 2.2e-16. (Right panel) Close-up of the agreement between experiment and model for contacts with probabilities less than 6%, which are not used as constraints in our modeling procedure. In this range, Pearson’s correlation is 0.907 with p-value = 0.004867. (C) The agreement between NE association frequencies from lamina-DamID experiments and the model population. This figure is plotted in the same way as (B). The structure population well reproduces the input frequencies derived from lamina-DamID data, with a Pearson’s correlation of 0.95 and p-value < 2.2e-16. (D) Comparison of experimental and model lamina-DamID frequencies on chrX. The top panel shows the input frequencies derived from the lamina-DamID signal, the middle panel shows the fraction of domains located at the NE in the structure population obtained by Hi-C and lamina-DamID data integration, and the bottom panel shows the fractions obtained in our control structure population generated using only Hi-C data.

Another quality measure for our models is how well we can predict the frequencies of chromatin interactions that were not included as constraints in the optimization. In our models, we did not impose constraints for any pair of TADs whose contact probability was lower than *a_ij_*=0.06. Very low contact probabilities are expected to contain a higher fraction of experimental noise. Such pairs include ~99.99% of all inter-chromosome and inter-chromosome arm interactions. However, our structure population is capable of predicting the missing data (**Fig. 2B** right panel). Many of the low-frequency contacts are formed as a consequence of imposing more significant interactions (with contact probabilities *a_ij_*>0.06), and their correct prediction is a good indicator of the model quality.

#### Reproducing the lamina-DamID binding frequency

Lamina-DamID experiments identify the probability that a locus is associated with the NE (more precisely, with the lamina protein located at the NE). We first assess the consistency between our structure population and the lamina-DamID experiment (a TAD domain-NE contact is defined when the domain surface is less than 50 nm from the NE). The association probabilities are in excellent agreement, with a Pearson’s correlation of 0.95 (**Figs. 2C, 2D** and **S1A**). Recalling that the TADs of Drosophila are divided into four functional classes, we find that TADs in the “Active” class are less frequently in contact with the NE than those from the other three classes (HP1, PcG, and Null) (Fig. S1A). This result agrees with prior observations in the literature that the genes interacting with lamina are usually transcriptionally silent and lack active histone marks [1]. The control population generated using only Hi-C data also shows good (albeit substantially lower) correlations between its NE association probabilities and the lamina-DamID experiments (Pearson’s correlation is 0.64, with p-value < 2.2e−16) (**Figs. 2D** and **S1B**). This relatively high correlation value in the control population shows a strong consistency between the Hi-C based models with the independent lamina-DamID data and confirms the generally good quality of our Hi-C based structure modeling.

#### Agreement with FISH experiments

Our genome structures also predict well the NE association frequencies observed by independent FISH mapping of 11 different genomic loci [1]. The Spearman’s rank correlation coefficient between experiment and model is 0.642 for these loci, with a significant p-value = 0.03312 (**Fig. S2A**). The corresponding correlation with the control structure population is substantially lower (Spearman’s rank correlation coefficient = 0.376 with p-value = 0.2542) (**Fig. S2B**), demonstrating the benefit of data integration to generate more accurate genome structures.

#### Presence of chromosome arm territories

Chromosome territories have been observed directly in higher eukaryotes, including mammalian cells [49, 50]. In Drosophila, chromosome territories can be inferred from the fact that Hi-C contact frequencies between chromatin regions in the same chromosome arms are substantially higher than those between chromosome arms [7, 8]. Previous 4C experiments on larval brain tissue confirm the limited nature of interactions between genes on different chromosome arms [41]. FISH experiments have also suggested chromosome territories in Drosophila [40]. In our models, we analyze the formation of chromosome territories by calculating a territory index (**TI**), which measures the extent of chromosome mixing [24]. To calculate **TI** in each structure, first we define the spanning volume of each chromosome, which is the surface convex hull of all its domain positions [24]. **TI** is then defined as the percentage of all domains occupying the chromosome spanning volume of the target chromosome (**Suppl. Methods C.2**). By definition, the maximum **TI** value of 1 indicates that the chromosome’s spanning volume is exclusively occupied by its own domains, and therefore experiences limited chromosome mixing. When considering domains from homologue chromosome copies, the territorial index ranges between 0.96 and 1.0 for all the chromosome arms (**Fig. S3A** and **Suppl. Table 2**). When separating the homologue chromosomes, however, the TI values range between 0.62 and 1.0 for the larger chromosome arms (**Fig. S3B**), suggesting that homologue chromosome pairs share almost the same territory due to strong homologue pairing.

#### Residual polarized organization

In a polarized genome organization, each chromosome occupies an elongated territory with the centromere at one nuclear pole and telomeres on the opposite side of the nucleus. Such an organization, called Rabl, typically occurs after mitosis and has been observed in a variety of plants [23], yeast, and both polytene and non-polytene Drosophila nuclei; it is also common in Drosophila embryos [27, 42, 43]. In the majority of our genome structures (67.4%, **Suppl. Methods C.3**), more than half of the chromosomes arms (chr2L, chr2R,chr3L, chr3R and chrX) are organized with their centromeres and telomeres located in opposite nuclear hemispheres (**Figs. S4B, C, and D**). This organization is also apparent when calculating the localization probabilities of chromosomes, which are highest for the telomeres in a region near the NE opposite to their respective centromeres (**Figs. 3A and B**). Taken together, these results suggest that interphase chromosomes retain some features of Rabl organization.

**Figure 3.**
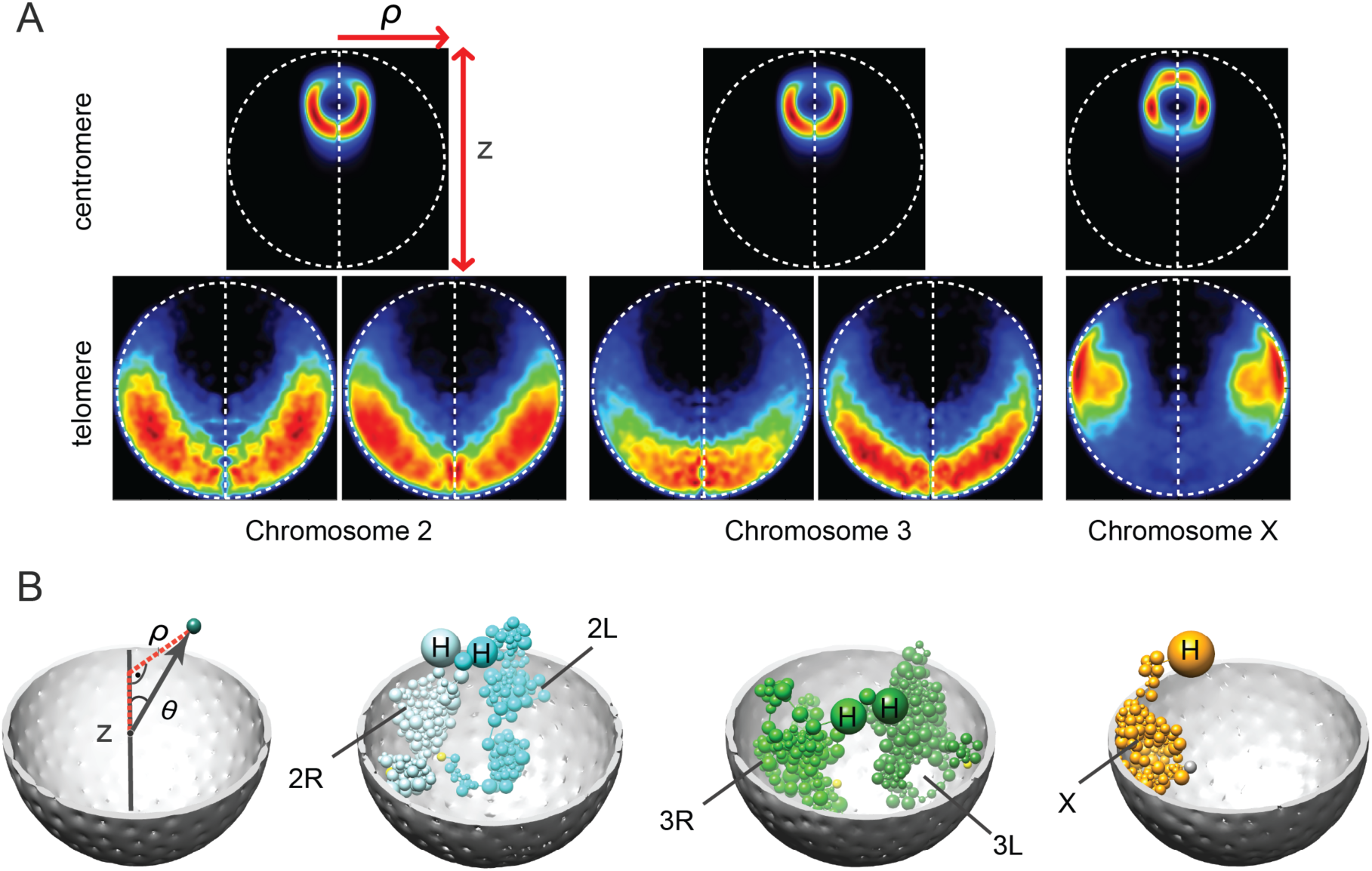
Residual polarized organization. (A) Projected localization probability densities (LPDs) of centromeres and peri-telomeric sequences for all chromosome arms calculated from the structure population. Probability densities are determined with respect to two principle axes of the nuclear architecture. The z-axis connects the center of the nucleolus with the origin at the nuclear center. The radial axis defines the distance of a point from the central z-axis (shown in the left panel in B). The left half of the projected localization density plot mirrors the right half for visual convenience. (B) Illustration of the genome organization for different chromosome arms in one genome structure.

#### Nuclear colocalization of Hox gene clusters

In Drosophila, the two PcG-regulated Hox gene clusters (Antennapedia complex and Bithorax complex) tend to co-localize in the head of 10-11 stage embryos [51], despite being separated by 10Mb in sequence on chromosome 3 (**Fig. 1B**). To test their spatial colocalization in our models, we calculate the pairwise spatial distances between the two gene clusters in every structure of the population (**Suppl. Methods C.4**). As a random control, we also calculate the pairwise distances between 30 pairs of gene clusters that only contain repressive TADs and share similar chromatin features in order to mimic the PcG-regulated Hox genes. In this control group each pair of gene clusters contains the same number of repressive domains, and are separated by the same sequence distance, as the pair of Hox gene clusters (**Suppl. Methods C.4**). We found that the Hox gene clusters are colocated in about 4.1% of structures in the population, a substantially higher rate than that observed in the control groups (median value 1.18%). Only 3 pairs of clusters among the 30 control groups are more frequently colocated than the Hox gene clusters (**Fig. S5** bottom panel). One of the three shows interactions between the pericentromeric regions and the Null domains. The other two pairs of gene clusters are brought together by nearby active domains, which form frequent interactions. Interestingly, our model does not impose contact constraints between the Hox gene clusters, because their contact probability was below 0.06 (**Fig. S5** top panel). Therefore, these results support the predictive power of our model.

#### White gene localizing near pericentromatic heterochromatin

Position-effect variegation (PEV) is a process whereby a euchromatic gene is deactivated through an abnormal juxtaposition with heterochromatin, due to chromosome rearrangements or transpositions. PEV has been intensively studied for the Drosophila *white* gene [52, 53], which is on the distal end of chromosome X and separated by more than 19 Mb from the pericentromeric heterochromatin region (**Fig. 1B**). A chromosome inversion can insert the *white* gene in sequence next to the pericentromeric heterochromatin, which leads to its repression. Hence, such chromosomal rearrangement may be favored if the *white* gene has an increased chance of being in spatial proximity to the heterochromatin. However, technical limitations prevent us from directly measuring contacts between the white gene and heterochromatin with Hi-C experiments. Using our structure population, we can measure how often the *white* gene is located close to the pericentromeric heterochromatin of chromosome X. As a control set, we took the four domains that are located at equivalent sequence distances to the heterochromatin regions on chromosomes 2 and 3.

Interestingly, the spatial distance between the *white* gene and the X chromosome heterochromatin is significantly smaller than the corresponding distances of the control groups (one-tailed Welch’s two sample t-test, p-value < 2.2e−16) (**Fig. S6A**). Although it is unlikely for distal loci to come together in 3D, we found that in ~1.3% of structures the *white* gene and the heterochromatin were positioned (within a distance of 200 nm) (**Fig. S6B**). This frequency is nine times larger than the colocalization frequency in the control sets (0.14% of structures). Therefore, our models reveal an increased propensity for the white gene to be located near the pericentromeric heterochromatin, compared to equivalent sites on other chromosomes. This result suggests that spatial proximity facilitates the occurrence of this translocation in living cells.

### Different chromosome domains have different preferred locations in the nucleus

The evidence listed above demonstrates the consistency of our models with experimental data and known properties of the Drosophila genome organization. Next, we describe new findings on the nuclear architecture and its functional significance based on our analysis of the model structure population.

#### Nucleolus and heterochromatin positioning

The nucleolus is a subnuclear structure linked to the assembly of ribosomal subunits. It is formed by nucleolar organizer chromatin regions (NOR), which contain the ribosomal DNA (rDNA) and are located close to the pericentromeric heterochromatin of chromosome X [45]. Our model allows the nucleolus to freely explore the nuclear space. However, its most likely radial position is between the center and periphery of the nucleus (**Figs. 4A** left panel and **S4A**). The large bodies of heterochromatin of each chromosome often enclose the nucleolus (**Fig. 4A**).

**Figure 4.**
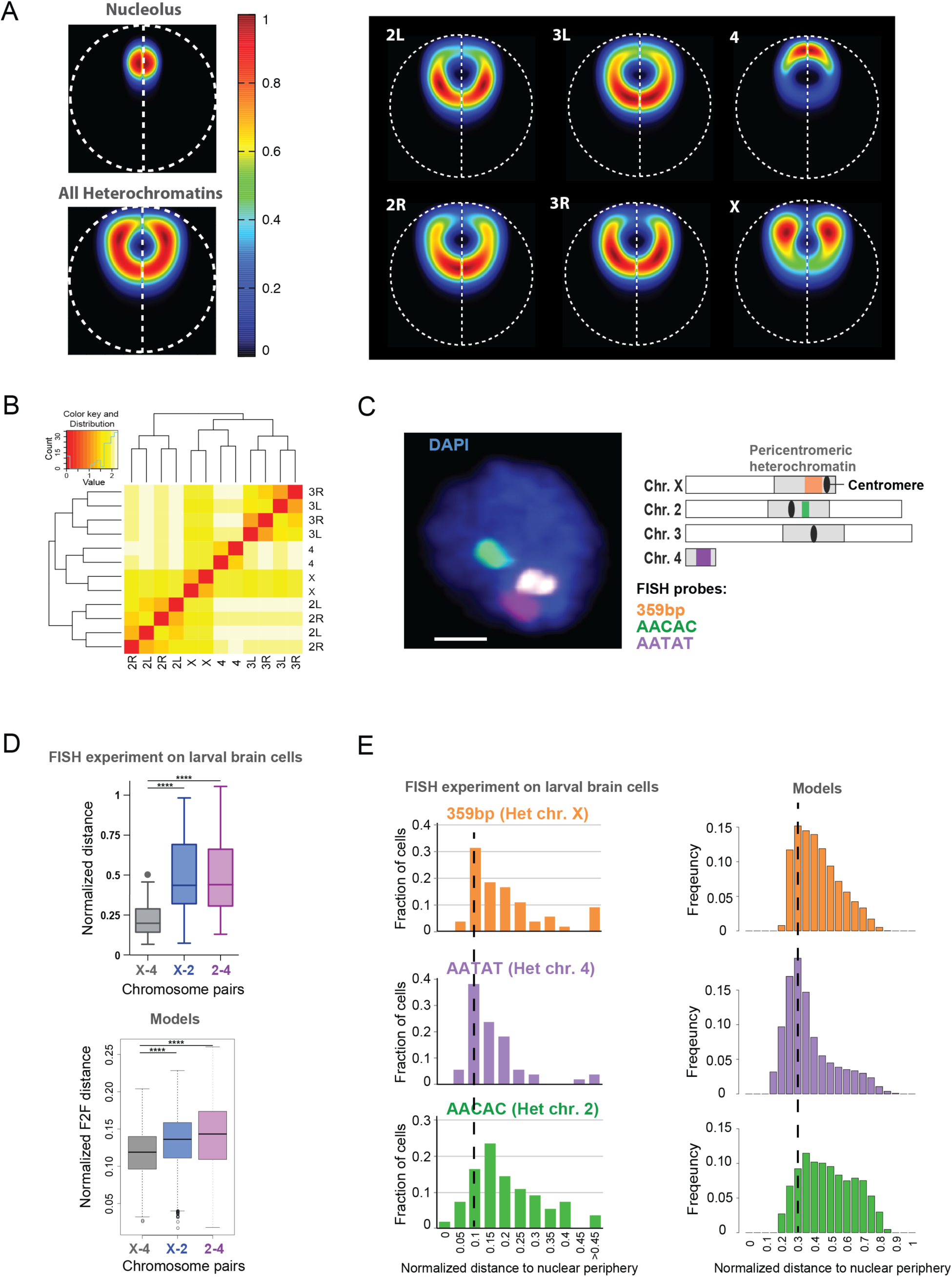
Heterochromatin and nucleolus positions. (A) (Left panel) LPD plots of the nucleolus and all pericentromeric heterochromatin regions in the model. On average, the nucleolus occupies an intermediate position between the center and the periphery, and is surrounded by pericentromeric heterochromatins. (Right panel) LPD plots for pericentromeric heterochromatin of different chromosome arms. They all exhibit different preferred locations. Those of chr4 and chrX are significantly more peripheral than the others. (B) Clustering of pericentromeric heterochromatin regions based on their averaged surface-to-surface distances. Heterochromatin domains of arms from the same chromosome naturally show preferred clustering. Heterochromatin domains from chr4 and chrX are usually closer to each other than to those from other chromosomes. (C) (Left panel) FISH signals in larval brain cells. The image shows the middle Z-stack of a representative nucleus. Scale bar = 1 μm. (Right panel) Schematic of the position of FISH probes used for this study, relative to the pericentromeric regions of each chromosome (chrX,chr2,chr4). (D) (Top panel) The positions of heterochromatic satellites from different chromosomes relative to each other, as measured by FISH experiments on larva brain cells. ****p-value < 0.0001 by paired t-test, N = 55 cells. (Bottom panel) Pairwise distances (surface-to-surface distance normalized by the diameter of the nucleus) between the heterochromatin domains as measured in the model. Similar to the data in vivo, the distance between the heterochromatin domains of chrX and chr4 is significantly smaller than the distance between the other two pairs, according to paired t-tests (p-value < 2.2e-16). (E) (Left panel) Positions of heterochromatic satellites from different chromosomes relative to the nuclear periphery, obtained from FISH experiments on larva brain cells. The heterochromatic satellites on chrX and chr4 are closer to the NE than those of chr2. (Right panel) A similar plot generated from the structure population shows good agreement with the FISH experiments.

Importantly, we validated this model prediction *in vivo*, using Drosophila Kc cells (**Fig. S7**). Immunofluorescence analysis of nucleoli and pericentromeric heterochromatin confirms that the average distance between the center of the nucleolus and the nuclear periphery is less than half of the nuclear radius (**Fig. S7B**). Interestingly, the nucleolus is positioned close to the nuclear periphery in 68% of cells, and close to the center of the nucleus in the remaining cells, revealing a bimodal distribution (**Figs. S7A, C**). In most cells, pericentromeric heterochromatin partially encloses the nucleolus (**Fig. S7A**). Interestingly, our model predicts certain location preferences for the heterochromatin of individual chromosomes. The heterochromatin regions of chromosomes 4 and X are usually positioned close to each other (**Fig. 4B and Supp. Methods C.5**), and both are more peripheral in the nucleus than the heterochromatin regions of chromosomes 2 and 3 (**Fig. 4A** right panel). The heterochromatin of chromosome 4 appears to be often positioned between the nucleolus and the NE (**Figs. 4A** right panel and **S4A**). We reason that the metacentric chromosomes 2 and 3 are roughly double the size of the acrocentric chromosome X, and therefore spread out more towards the interior of the nucleus. Notably, we confirmed these predictions using FISH staining of heterochromatic repeated sequences (satellites) in Drosophila cells of larval brains. As shown in **Fig. 4C**, the satellite repeats of chromosomes X and 4 are more often closer to each other than those of chromosomes X and 2, or 2 and 4 (**Fig. 4D** top panel), in agreement with our models (**Fig. 4D** bottom panel). Further, the satellite repeats of chromosomes X and 4 are more often closer to the nuclear periphery than those of chromosome 2 (**Fig. 4E** left panel), which also confirms our findings in the model structure population (**Fig. 4E** right panel). Together, these *in vivo* data support our model, and suggest that the predicted chromosome organization is not limited to embryonic cells.

#### Localization of all euchromatin domains

When plotting the average radial position for every euchromatic TAD (**Fig. 5A**) we observe that the arms near the pericentromeric heterochromatin regions are preferentially positioned in the nuclear interior, while euchromatic regions at the telomeric ends prefer the periphery. This preference is also seen for chromosome 4, despite its small size.

**Figure 5.**
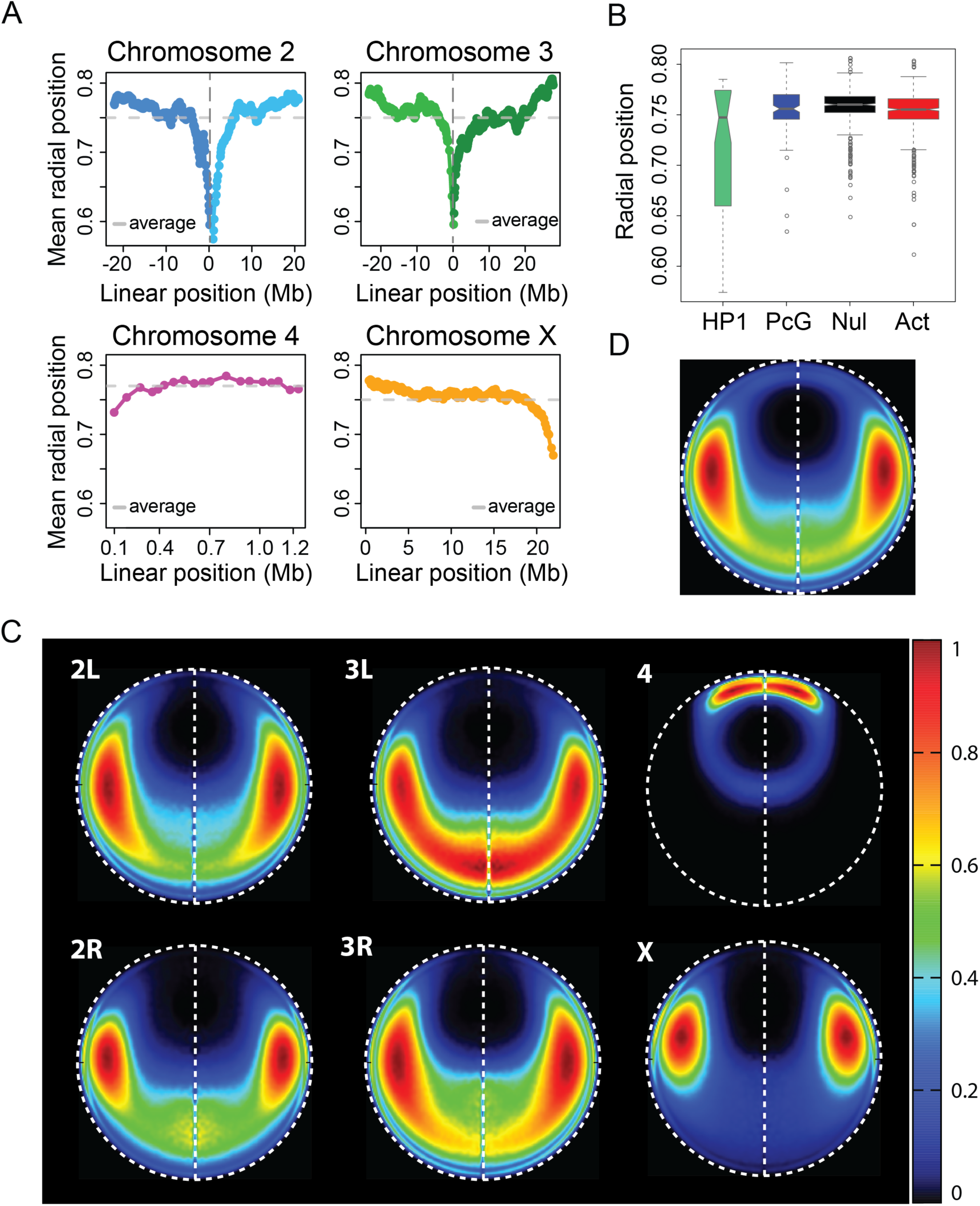
Localization of euchromatin domains in the structure population. (A) The average radial position for each euchromatin domain, plotted by position along its chromosome. The 0 location along the x-axis of chr2 represents the euchromatin region closest to the centromere, with 2L domains on the left and 2R domains on the right. Chr3 domains are plotted with the same coordinate system as chr2. The domains of chr4 are plotted from left to right, while the domains of chrX are plotted from right to left; this convention follows the schematics in Figure 1. Centromeric regions and pericentromeric heterochromatin regions are not shown in this figure. The domains near pericentromeric regions are closer to the nuclear center on average, while the domains near telomeric ends are preferentially close to the nuclear periphery. (B) The average radial positions of each domain, grouped by epigenetic class. (C) LPD plots of all euchromatin domains from each chromosome arm in nuclear space. (D) LPD plot of all euchromatin domains.

Euchromatic regions are either active or repressed, and can be divided into 4 classes based on their epigenetic profiles: Null, Active, Polycomb-Group (PcG), and HP1 [8] (**Suppl. Table 3**). The TADs of the Null, Active, and PcG classes have similar average radial positions (**Fig. 5B**). The average radial positions of the HP1 TADs have larger variance. The pericentromeric HP1 TADs (excluding all TADs on chr4) are found near the nuclear interior substantially more often than non-pericentromeric HP1 TADs.

Based on our model structures, we can create localization probability density plots (LPD) for the euchromatic regions of different chromosomes (**Fig. 5C**). The chromosome with the most distinct location preference is number 4, whose euchromatic regions reside very close to the NE. In contrast, a large part of chromosome 3L is located on the side of the NE opposite to chromosome 4 along the central axis, coinciding with the line drawn between the centers of the nucleus and nucleolus (vertical dashed line in **Fig. 5C**). Chromosome 2, on the other hand, prefers to avoid the central axis. The right and left arms have similar location preferences. The location distributions of chromosomes 2 and 3 are qualitatively similar, but chromosome 3 is more likely to be found close to the central axis. Chromosome X resides fairly close to the nucleolus, around the midpoint of the central axis, and is considerably less dispersed than the arms of chromosomes 2 and 3.

### Analysis of homologous pairing

#### Distances between homologous pairs vary along the chromosome

*D. melanogaster* shows somatic homologous chromosome pairing in interphase nuclei [31–33, 44]. Moreover, the paired chromosomes touch only at a few specific interstitial sites [31]. In our structures, we define a domain as being paired if the surface-to-surface distance between the two homologs is less than 200nm (**Fig. 6A**). Interestingly, the pairing frequencies of homologous domains show distinct and reproducible variation along the chromosomes (**Fig. 6B** left panel). The active class shows smallest homologous pairing frequency for each chromosome (**Fig. 6B** right panel). During the optimization, all pairs of homologue TAD copies are subject to a generic upper bound constraint, which limits their maximum separation to 4 times the TAD diameter. Even though this constraint is the same for all domains, it is noteworthy that in the optimized structures, certain pairs of homologue TADs consistently have small average separations while others consistently have separations close to the upper bound. Hence, this distance variation is TAD-specific and highly reproducible in independently calculated structure populations (**Fig. 6C**). This effect is an indirect consequence of the genome-wide Hi-C and lamina-DamID constraints imposed on the structures.

**Figure 6.**
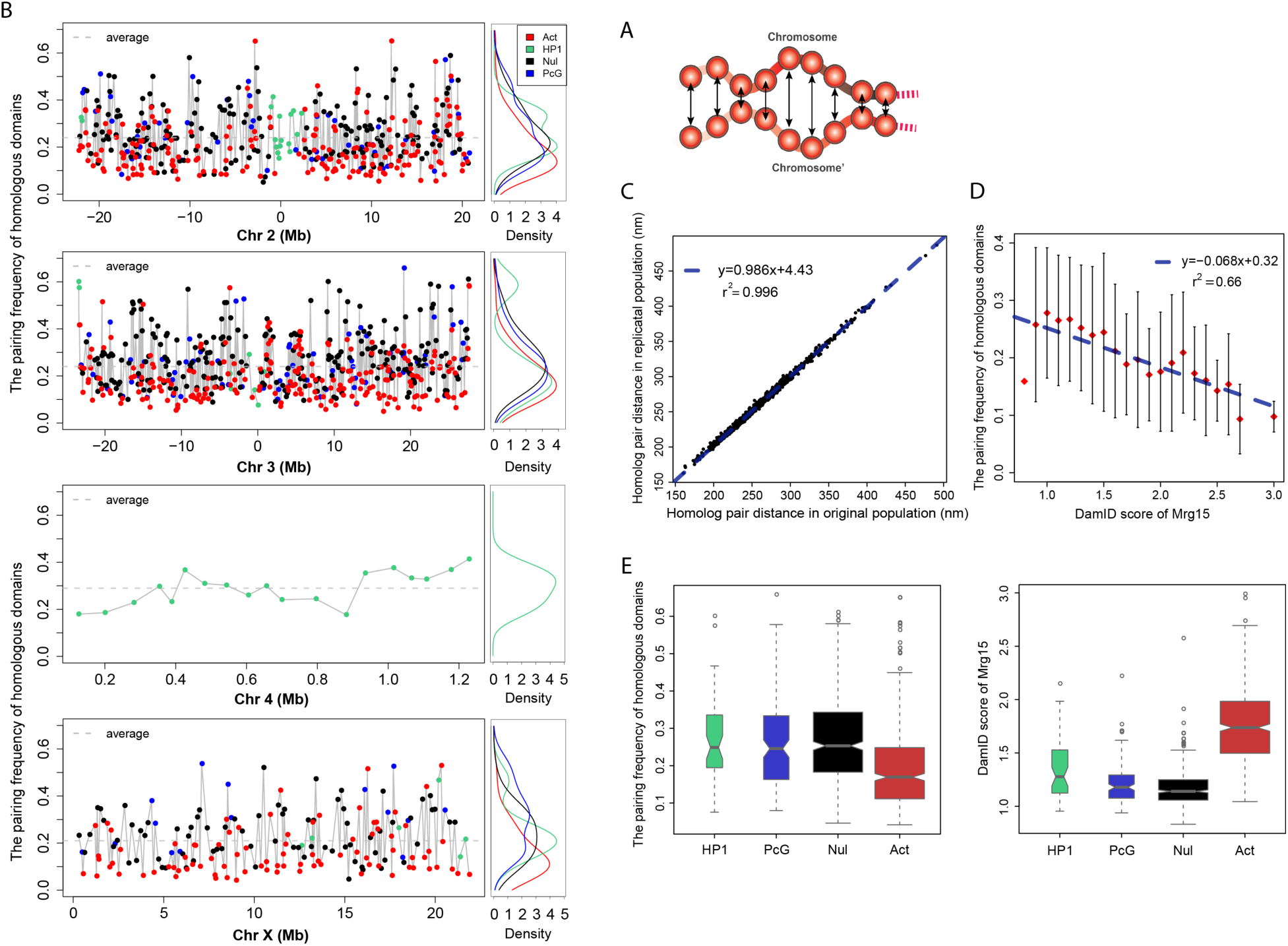
Analysis of homologous pairing. (A) Schematic view of surface-to-surface distances between homologous domains. Different domains exhibit different degrees of homologue pairing. (B) (Left panel) Pairing frequency for each euchromatin domain, plotted by chromosome. We define a domain as being paired in a structure if the surface-to-surface distance between the two homologs is less than 200nm. The X-axes are the same as the plots in Fig. 5A. The domains are colored by their epigenetic classes: Green-HP1, Blue-PcG, BlackNull, and Red-Active. (Right panel) Density plot of the domain pairing frequencies, grouped by epigenetic class. The active class has the smallest mean homologous pairing frequency for each chromosome. (C) Reproducibility of the average homologue distances between two independently generated structure populations. The Pearson’s correlation between them is 0.998, with p-value < 2.2e-16. (D) There is a negative correlation between the pairing frequencies of homologous domains and their Mrg15 enrichment. The Mrg15 scores range from 0. 8 to 3.0, and are divided into 21 equal bins. The corresponding pairing frequencies from our models in a given Mrg15 bin are summarized as a mean and variance, and the latter is displayed as an error bar. The blue dotted line is the linear regression between the average pairing frequency in each bin and the midpoint Mrg15 enrichment value of the bin. The Pearson’s correlation between them is −0.8126852, with p-value = 7.591e-06. (E) (Left panel) Pairing frequencies of homologous domains, grouped by epigenetic class. (Right panel) Enrichments of Mrg15 binding, grouped by epigenetic class. Active domains are generally more enriched with Mrg15, and have lower pairing frequencies, than the other three repressive classes.

The consistency of this pairing behavior raises the question of why certain regions attain higher levels of pairing. One clue is that we find a small but significant correlation between pairing frequency and the location of the TAD in the nucleus. Pearson’s correlation between the frequency of pairing and the frequency of being in proximity to the NE is 0.34 (p-value < 2.2e−16) (a TAD-NE contact is defined when its domain surface is is less than 50nm from the NE). We hypothesize that genomic regions that are often positioned near the NE may be more restricted in their movements, which may facilitate homolog pairing. We also investigated whether the local crowdedness around the domains could influence the spatial distances between homologues, and found that in the majority of structures the local crowdedness is not different between paired domains and unpaired domains (**Supp. Methods C.6**).

#### Mrg15 is enriched in active domains and depleted in repressive domains

Several proteins have been reported to affect somatic homolog pairing in Drosophila [32, 33, 44]. Among them is Mrg15, which binds to chromatin and recruits CAP-H2 protein to mediate homolog unpairing [44]. Interestingly, we find an anticorrelation between Mrg15 binding enrichment in a domain and a domain’s homologous pairing frequency, even though this information is not imposed as an input constraint in our models (**Fig. 6D**). The higher the Mrg15 enrichment signal in a domain, the lower the fraction of paired homologues in the structure population (**Fig. 6D**). Pearson’s correlation coefficient between the binned Mrg15 binding signal and the averaged frequency of homologous pairing for each bin is −0.81, with p-value = 7.59e-06 (**Fig. 6D**). In the control model (using only Hi-C data), the Pearson’s correlation coefficient between them is −0.697 with p-value = 0.000446. We also divided the domains into three subsets based on their Mrg15 scores. The average pairing frequency for domains enriched with Mrg15 is significantly less than that for domains with lower Mrg15 scores (one-tailed Mann-Whitney U test, p-value < 2.2e−16) (**Fig. S8A**).

Among the four TAD classes, “Active” domains are generally more enriched with Mrg15-binding sites (**Fig. 6E** right panel). Appropriately, we observe that transcriptionally active domains have a lower pairing frequency than the three repressive classes (**Fig. 6E** left panel). The most intuitive explanation is that a loose pairing makes an active domain more accessible to regulatory factors. PcG domains, which are enriched with polycomb group proteins, show higher levels of homologous pairing in our models than the active domains (one-tailed Welch’s two sample t-test, p-value = 2.09e−9). Therefore, our structure population supports the notion that PcG domains form tight pairs to enhance gene silencing (reviewed in ref. [26]).

While active domains generally have low frequencies of homologous pairing, our models also have some clear and reproducible counterexamples of active domains with extremely high frequencies of homologous pairing (the specific TADs with this behavior are reproducible in independently generated structure populations) (**Fig. 6C**). Therefore, we divided the active domains into two subclasses, labeled “active-tight” and “active-loose”. Interestingly, domains in the active-loose subclass have significantly higher Mrg15 enrichment than domains in the active-tight subclass (one-tailed Mann-Whitney U test, p-value = 0.03436) (**Fig. S8B**). It is interesting that our model further supports a role for Mrg15 in disrupting homologue pairing, even though the structures were generated without any locus-specific constraints on the separation of homologous domains. Importantly, the anticorrelation between homologue pairing frequency and Mrg15 binding signal further increases when lamina-DamID data is integrated in the model, which indicates that data integration helps generate more accurate genome structures.

#### Active-tight domains show higher transcriptional efficiency

Interestingly, we found significant functional differences between active-loose and active-tight domain subclasses. Active-tight domains contain more genes (**Fig. 7A**). Surprisingly, the active-tight subclass shows significantly lower binding levels of the TATA-binding protein (TBP) and RNA polymerase II, as well as lower H3K4me2 signals (one-tailed Mann-Whitney U test, p-values are 1.17e−04, 2.95e−03 and 5.19e−04 respectively) (**Fig. 7B**). However, the gene expression levels in the two subclasses are comparable, despite the significantly smaller amount of bound RNA Pol-II transcription machinery in the active-tight subclass. This observation suggests that homologue pairing of active alleles might improve transcription efficiency even at lower concentration of transcription factors.

**Figure 7.**
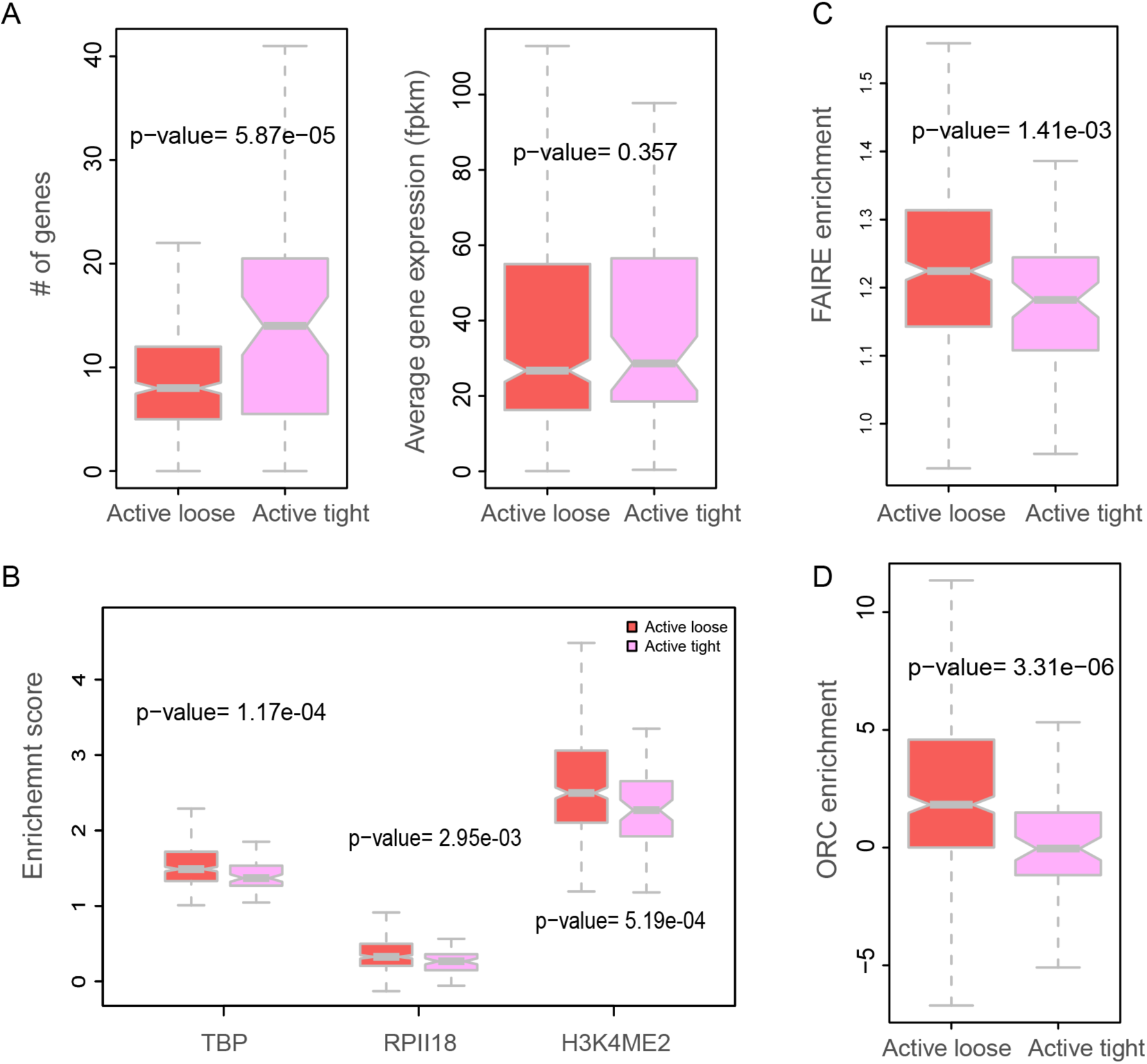
Transcriptional efficiency and DNA replication timing for genes in two subclasses of the Active domains. (A) Domains in the “active-loose” subclass have lower frequencies of homologue-pairing than those in the “active-tight” subclass (Supp. Methods C.6). The active-tight subclass includes 71 domains, and the active-loose sub-class includes 423 domains. All the statistical tests are performed using one-tailed Mann-Whitney U test. (Left panel) Domains in the active-tight subclass contain significantly more genes than domains in the active-loose subclass. (Right panel) Genes in both sub-classes have similar average expression values. (B) TBP (TATA binding protein), PolII binding signal and H3K4me2 signals are more enriched in domains of the active-loose subclass. (C) FAIRE signal is significantly stronger in domains of the active-loose subclass. (D) ORC is significantly more enriched in domains of the active-loose subclass.

#### Active-tight domains tend to be late-replicating

FAIRE (Formaldehyde-Assisted Isolation of Regulatory Elements) is a biochemical method to identify nucleosome-depleted regions in the genome. It has been shown that these DNA sequences overlap with active regulatory sites and DNaseI hypersensitive sites [54]. Active-loose domains are significantly enriched with in the FAIRE signal compared to domains in the active-tight subclass (**Fig. 7C**). This indicates that chromatin in the active-loose domains is more depleted of nucleosomes, and hence these domains contain a higher density of regulatory chromatin complexes. In Drosophila, the organization of nucleosomes plays an important role in determining origin recognition complex (ORC) binding sites [55]. The difference in FAIRE enrichment leads us to investigate DNA replication timing during interphase for the different classes. The Active domains are generally more enriched with ORC than the other three types, with significant p-values (one-tailed Mann-Whitney U test, p-values are 3.259e−15, 0.001715 and 0.01837 for NULL, HP1 and PCG respectively), indicating that DNA replication is often initiated in the chromatin of the active class (**Fig. S9B**). Strikingly, we discovered that ORC-binding regions are much more frequent in active-loose domains than in active-tight domains (one-tailed Mann-Whitney U test, p-value=1.54e−4) (**Fig. 7D**), supporting the model that chromatin in the active-loose subclass replicates significantly earlier (i.e., in early S-phase as opposed to late S-phase).

## Discussion

It has become increasingly clear that a chromosome’s folding pattern and nuclear location have far-reaching impacts on the regulation of gene expression and other genome functions. Therefore, a thorough understanding of a genome’s function entails detailed knowledge about its spatial organization. A wide range of complementary technologies exists to provide such information. For instance, genome-wide ligation assays provide critical information about chromatin-chromatin interactions, lamina-DamID experiments reveal the propensity of a given locus to be located close to the NE, and 3D imaging technologies can reveal the spatial locations of individual loci in single cells. However, many computational models of genome structures rely on a single data type, such as Hi-C, which limits their accuracy. Integrating complementary data types increases the accuracy and coverage of genome structure models, and also provides a way to cross-validate the consistency of data obtained from complementary technologies. Thus, a major and vital challenge of computational biology is to develop hybrid methods that can systematically integrate data obtained from different technologies to generate structural maps of the nucleome (e.g., as this study integrates Hi-C and lamina-DamID).

In this paper, we present a computational platform that can systematically integrate experimental data obtained from different technologies to map the 3D structures of entire genomes. Our probabilistic approach explicitly models the variability of genome structures between cells by simultaneously deconvolving data from Hi-C and lamina-DamID experiments into a model population of distinct diploid 3D genome structures. Our models therefore incorporate the stochastic nature of chromosome conformations, and allow a detailed analysis of alternative chromatin structure states.

Our method can be applied to genomes of any organism, including mammalian genomes. As a proof of principle, we mapped the structure of the *D. melanogaster* genome in interphase nuclei. We demonstrated that our method produces an ensemble of genome structures whose chromatin contacts are statistically consistent with Hi-C data while also reproducing the likelihoods of chromatin loci being close to the NE derived from lamina-DamID experiments.

The ensemble of model structures has strong predictive power for structural features not directly visible in the initial data sets. We observed that, in embryonic cells, chromosomes 2 and 3 are often organized with their centromeres and telomeres located in opposite hemispheres of the nucleus. In addition, each chromosome pair occupies a distinct territory in our models. Our structures also predicted correctly a relatively high colocalization probability between the two PcG-regulated Hox gene clusters, even though no contact constraints were imposed between these genes when the model structures were generated.

Due to technical limitations, no Hi-C measurements are available to confirm interactions of heterochromatin regions. However, using our 3D model structures, we can analyze the positions of chromatin loci with respect to heterochromatin regions. For instance, our model shows a high preference for the *white* gene on chromosome X to be positioned close to pericentromeric heterochromatin in comparison to similar gene locations on other chromosomes, thus facilitating the *white* gene’s translocation next to heterochromatin. Our analysis also reveals distinct differences between some chromosomes in terms of heterochromatin localization probabilities. For example, pericentromeric heterochromatin of chromosomes X and 4 are more proximal to each other than to pericentromeric heterochromatin of chromosomes 2 and 3. The preferred euchromatin locations of chromosome 4 are also distinctly different from those of the other chromosomes.

We also make intriguing observations about homologous pairing that cannot be directly observed in the original Hi-C or lamina-DamID data. In our models, the tendency for domains to pair varies a great deal along the chromosome, which confirms the idea that pairing initiates from several distinct loci and spreads to neighboring regions. The observed pairing tendency of the domains is highly reproducible over several independent simulations, and also correlates with distinct functional features of the domains. We investigated why certain domains are more frequently paired than others. Interestingly, there is an anti-correlation between pairing frequency and the enrichment in Mrg15 protein binding, which is known to affect somatic chromosome pairing in Drosophila. This information was not explicitly included in the modeling process. The pairing frequencies of homologous domains also differ between those containing active or repressed chromatin. Active domains generally have a lower frequency of chromosome pairing than repressed domains such as those enriched in the polycomb groups (PcG) of proteins. However, we also identified some active domains that break this pattern, with extremely high rates of chromosome pairing across many independent simulations. Interestingly, when we compare these outlier active domains with the more common type of active domain having low pairing frequencies, the former have substantially lower levels of Mrg15 binding signals, later DNA replication timing, and lower FAIRE signals. These attributes are similar in other regions with high pairing frequencies.

Homologous pairing has been studied for years, and it has been found to play a large role in gene regulation. Transvection is a phenomenon whereby gene expression is modulated by the physical pairing of homologous loci. A case study showed that more transcripts are produced when both alleles of the gene Ubx are paired than when they are spatially separated [28]. A possible explanation is that each gene copy can be activated by both its own and the other copy’s enhancer [26]. Interestingly, when we compare actively transcribed genes in chromatin regions with very high or very low levels of homologous pairing, the former show significantly lower signals in RNAPII and TATA protein binding, but at the same time similar levels of transcripts. This observation indicates that a more efficient transcription of genes occurs when pairing is frequent. Our model also shows that regions with looser homologue pairing initiate replication earlier than regions with tighter homologue pairing.

## Conclusions

In this study, we address one of the principal challenges of genome structure analysis: the development of a method that systematically integrates complementary data from different technologies to map the 3D organizations of genomes. Data from a single source, such as Hi-C or lamina-DamID experiment alone, cannot capture all aspects of a genome’s organization. Integrating multiple data types is therefore not just beneficial but necessary to enhance the accuracy and coverage of structural models. Furthermore, the detailed analysis of such structural models is a valuable complement to experimental studies, because it can provide new structural insights. For example, the 3D models can reveal the relative locations of specific chromatin regions in the nucleus which are not immediately visible in the initial data. In the future, genome structure modeling should rely on all available data, including live fluorescence and 3D FISH imaging, as well as Hi-C and lamina-DamID experiments from both large-scale single cell and ensemble technologies. This approach will permit an extremely detailed analysis of the genome’s structural features, at high resolution and fully consistent with all experimental findings. Our work is a first step towards this goal, in that it allows the integration of genome-wide Hi-C as well as lamina-DamID data for 3D genome structure analysis, and provides a robust computational framework for integrating structural constraints from other types of experiments.

## Materials and Methods

### General description

The population-based approach is a probabilistic framework to generate a large number of 3D genome structures (i.e., the structure population) whose chromatin domain contacts are statistically consistent with experimental Hi-C data and other spatial constraints derived from *a priori* knowledge and/or independent data types. Our model is a deconvolution of the ensemble-averaged Hi-C data, and the resulting structures can be considered the most likely representation of the true structure population over a population of cells, given all the available data. Our method distinguishes between interactions involving chromosome homologues, so it can generate structure populations representing entire diploid genomes. Further, because the generated population contains many different structural states, this approach can accommodate all experimentally observed chromatin interactions, including those that would be mutually exclusive for a single structure. Compared to our previous research, which introduce the population-based approach using Hi-C data alone, in this study we also integrate lamina-DamID data to generate an improved structure population.

### Chromosome representation

The nuclear architecture of Drosophila cells consists of the nuclear envelope (NE), the nucleolus, and eight individual chromosomes (the diploid pairs chr2, chr3, chr4 and chrX). Chr2 and chr3 each have two arms, labeled 2L-2R and 3L-3R, connected by centromeres (**Fig. 1B**).

Each chromosome contains three main regions: euchromatin, pericentromeric heterochromatin, and a centromere (**Fig. 1B**). Euchromatin regions in chromosome arms 2L, 2R, 3L, 3R, 4 and X are linearly partitioned into a total of 1169 well demarcated physical domains [8], which are represented as spheres in the model [22]. A domain sphere is characterized by two radii: (1) its hard (excluded volume) radius, which is estimated from the DNA sequence length and the nuclear occupancy of the genome; and (2) its soft (contact) radius which is twice the hard radius. A contact between two spheres is defined as an overlap between the spheres’ soft radii. This two-radius model allows for the possibility that chromatin can partially loop out of its bulk domain region to form contacts, while establishing a minimum genome occupancy in the nucleus. According to experimental data, the combined hard-core spheres of all euchromatin domains occupy around 12% of the nuclear volume. The total volume of heterochromatin is set to 1/27 of the nuclear volume. This figure is in agreement with estimates from microscopy images ([46] and **Fig. S7A**), which show the heterochromatin cluster to occupy roughly one third of the nuclear diameter. The heterochromatin regions of each chromosome are modeled as spheres occupying volumes proportional to 5.4: 11.0: 8.2: 8.2: 3.1: 20.0, according to the chromosome outlines depicted in **Fig. 1B** (these volumes are taken from the data shown in ref. [56]). For every chromosome, the centromere is modeled as a sphere with 5% the volume of its corresponding heterochromatin domain (or sum of two heterochromatin domains for chr2 and chr3).

The nuclear radius is set to 2 microns (μm) as suggested by fluorescence imaging experiments ([35, 46] and **Figs. 4C, S7A**). The nucleolus radius is set to 1/6 of the nuclear radius (**Fig. S7A**). Centromeres are clustered together and attached to the nucleolus [35]. Pericentromeric heterochromatin of chrX surrounds the rDNA cluster regions, so it lies in close proximity to the nucleolus. (**Suppl. Table 3** lists all domain radii in the model.)

All these units are represented by a total of 2359 spheres (**see Table 1**).

**Table 1:** Structural units of our *Drosophila melanogaster* genome model

| Genome component | Unit quantity | Number of spheres | Description |
| --- | --- | --- | --- |
| TAD | 1169 | 2338 | Euchromatin TADs |
| HET | 6 | 12 | Heterochromatin clusters on 2L, 2R, 3L, 3R, 4, X |
| CEN | 4 | 8 | Centromeres of chromosomes 2, 3, 4, and X |
| Nucleolus | 1 | 1 | Localization of nucleoli |

The outlines of the chromosomes are depicted in **Fig. 1B**. In the next section, we briefly describe the chromosome model and list all of the structural constraints that we imposed while optimizing the population.

### Probabilistic platform for data integration

Our method closely follows our recent population-based modeling framework [22]. However, we now generalize this framework to support the integration of lamina-DamID data with Hi-C data. The Hi-C data is contained in the ensemble contact probability matrix **A**, and the lamina-DamID data is contained in the ensemble chromatin-NE contact probability vector *E*.

We aim to generate a structure population **X** that maximizes the likelihood *P*(**A**,*E*|**X**). We introduce two latent variables **W** and **V**, which represent features of individual cells that aggregate into the ensemble information **A** and *E*, respectively. **W** = (*w_ijm_*)_2*N*×2*N*×*M*_ is the contact indicator tensor, which contains the missing information in the Hi-C data **A**: the presence or absence of contacts between all domain homologues, in each structure of the population ( *w_ijm_* = 1 indicates a contact between domain spheres *i* and *j* in structure *m*; *w_ijm_* = 0 otherwise). The second latent variable, **V** = (*v_im_*)_2*N×M*_, contains information whether each domain homologue is located near the NE, in each structure of the population (*v_im_* = 1 indicates that domain sphere *i* is near the NE in structure *m*; *v_im_* = 0 otherwise). Note that while these latent variables are indexed over domain homologues (lowercase indices *i, j*), which are independent spheres in the model, the ensemble datasets **A** and *E* in the formulas below are indexed over haploid domain identities observed in the experimental data (uppercase indices *I, J*). The maximum likelihood problem is then formally expressed as Eq. (1) and the expansion form is described as in Eq. (2).

Furthermore, *P*(**W,V**|**X**) can be expanded into a product of every contact indicator probability, i.e. 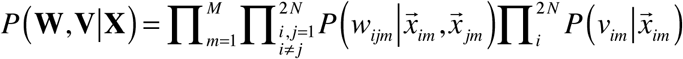 Then the term *P*(**A**|**W**) can be expanded as *P*(**A**|**W**) = **Π**_*I,J*_*P*(*a_IJ_*|*a′*_*IJ*_) where *a′*_*IJ*_ is the contact probability of the domain pair *I* and *J*, 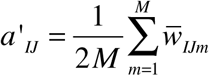. The projected contact tensor **W̅** = (*w̅_IJm_*)_*N*×*N*×*M*_ is derived from **W** by aggregating its diploid representation to the haploid counterpart.

Likewise, *P*(*E*|**V**) = **Π**_*I*_*P*(*e_I_*|*e′_I_*), where *e′_I_* is the probability for domain *I* to be near the NE. This is calculated as 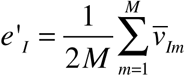. The term *v̅_Im_* is a matrix element of the projected matrix **V̅** = (*v̅_lm_*)_*N*×*M*_, and indicates how many domain *I* representations in structure *m* are near the NE; thus, its possible values are {0, 1, 2} when the diploid representation is projected to the haploid counterpart.

With these probabilistic models, we can maximize the log-likelihood log *P*(**A**, *E*, **W**, **V** | **X**), expressed as follows:

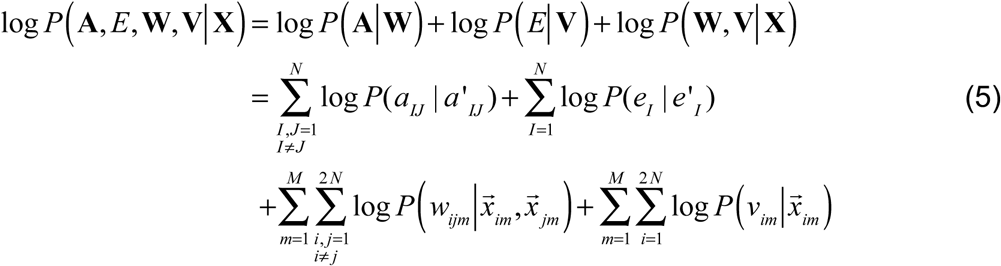

We assume that a pair of spheres (*i, j*) are in contact in structure *m* if and only if their center distance 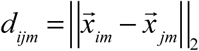 is between certain lower and upper bounds, *L*≤*d_ijm_*≤*U*. The lower bound is the sum of their hard radii, *L* = *R_i_* + *R_j_*, and the upper bound is the sum of their soft radii, *U* = 2(*R_i_* + *R_j_*). We modeled the probability of a contact between two domain spheres *i* and *j* as a variant of the rectified or truncated normal distribution, expressed as follows.

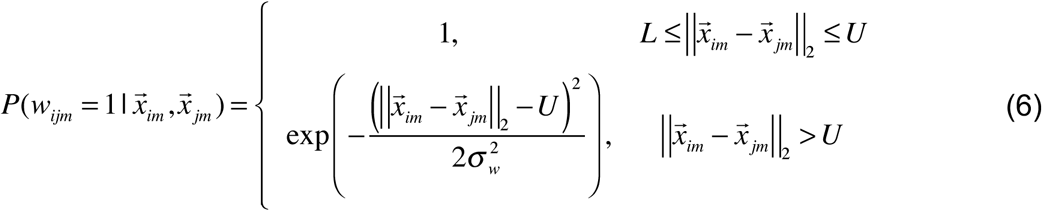

with very small variance, e.g. *σ_w_* → 0.

The probability for a domain to reside near the NE is described as

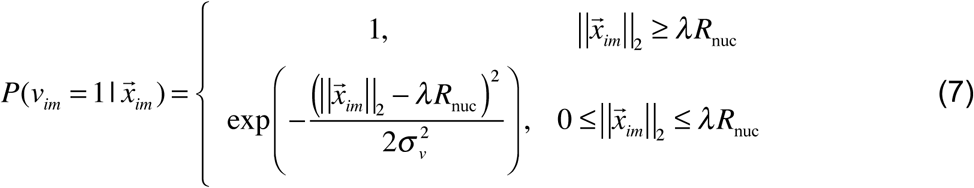

where *λ* = 0.975 to ensure that the enforced TAD is at the inside surface of the NE, and likewise *σ_v_* → 0.

### Additional spatial constraints for the Drosophila genome

In addition to the data from Hi-C and lamina-DamID experiments, we include the following additional information as spatial constraints:

1. *Nuclear volume constraint:* All 2359 spheres are constrained to lie completely inside a sphere with radius R_nuc_, i.e. 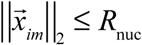. Without loss of generality, we use the origin (0,0,0) as the nuclear center, so 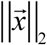 is the distance from the nuclear center.
2. *Excluded volume constraint:* The model prevents any overlapping between the 2359 spheres, as defined by their hard radius. For every pair of spheres *i* and *j* in every structure *m*, we enforce 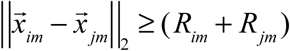.
3. *Homologue pairing constraint:* Based on experimental evidences, homologous chromosomes are somatically paired in Drosophila and so both copies of a gene are usually close to each other [30–33]. Therefore, we constraint the distance between 2 homologous domains to be less than an upper bound, which is four times the sum of their radii i.e. 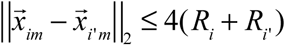.
4. *Consecutive TAD constraint:* To ensure chromosomal integrity, we apply an upper bound to the distance between two consecutive TAD domains, which is derived from the experimentally determined contact probability *a_ij_*. The upper bound distance is 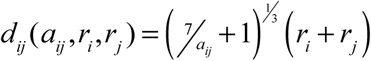. Note that *d_ij_* = 2(*r_i_* + *r_j_*) when *a_ij_* = 1.
5. Additional knowledge-based chromosome integrity *constraints:* The heterochromatic region of a given chromosome or chromosome arm forms a clustered subcompartment, so is represented by a single domain. No Hi-C data are available for the heterochromatic regions. To ensure chromosome integrity, the domains representing heterochromatic regions are always in contact with their adjacent TAD as well as with the centromeric domain. The constraint between the heterochromatin sphere and the adjacent TAD sphere *i* is 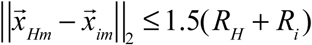. The constraint between the heterochromatin domain and the adjacent centromere sphere is 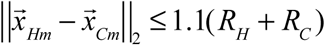, where 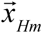 and 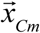 are the centers of the heterochromatin and centromere spheres, and *R_H_* and *R_C_* are the hard radii of the heterochromatin and centeromere spheres. Based on experimental evidence [35], all centromeres are in proximity to the nucleolus. Therefore, we constrain the centromere spheres to be close to the spherical volume representing the nucleolus, defined as 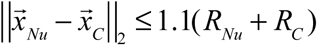 where *R_Nu_* is the radius of the nucleolus volume.

### Distance threshold method for estimating W and V

We adopt the distance threshold method introduced elsewhere [22] to estimate the distribution of contacts among the diploid genome across a population of structures. The distance threshold 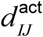 for each domain pair (*I,J*) is determined based on the empirical distribution of all distances between their homologous copies across all structures of the population. The procedure to determine a distance threshold for estimating an element of the projected contact indicator tensor, *w̅_IJm_*, is as follows. Let (*I, J*) be a domain pair (with homologues *i*, *i*’ and *j*, *j*’) and let their Hi-C contact probability *a_IJ_* > 0. We construct an empirical distribution of the pairwise domain distances between homologous copies of the domain pair (*I, J*). When *I* and *J* are domains from the same chromosome, we collect the distances *d_ijm_* and *d_i′j′m_*, in all model structures (*m*=1, 2,…, *M*), forming a set of 2*M* distances. When *I* and *J* are domains from different chromosomes, we collect the smallest 2 distances from the set of all possible distances {*d_ijm_*, *d_i′jm_*, *d_ij′m_*, *d_i′j′m_*,}, again for a total set of 2M distances. Next, the 2*M* distances are ranked in increasing order. The distance threshold, 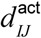, is defined as the distance value with the (2*M* · *a_IJ_*) th rank among the 2*M* sorted distances. Once all the distance thresholds are obtained, we populate the tensor **W̅** by counting how many of the pooled distances between (*I, J*) from structure *m* in the set of 2*M* distances that fall below the corresponding distance threshold. The structure optimization then assigns contacts to the pairs with shorter distance out of 4 possible pairs between homologue domains, for every *w_ijm_*. This procedure maximizes log*P*(**A, W|X**), which is composed of two items: log *P* (**W|X**) and log *P* (**A|W**). This is true for two reasons. (i) It assigns contacts only to domain pairs with short distances, maximizing log *P* (**W|X**). (ii) It uses the *2a_IJ_M*^th^-quantile of all 2*M* distances as the distance threshold to determine *w_ijm_*, which heuristically maximizes the first term 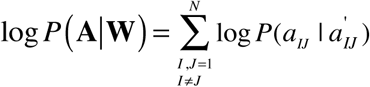 by making *a_Ij_* exactly equal to *a′_IJ_*.

We adapted this procedure to estimate the TAD-NE contact matrix **V** = (*v_im_*)_2*N×M*_. The distance threshold for every TAD is determined. Again we sort a set of 2M distances to the NE related to domain *I* in increasing order, and select the (2*M* ·*e_I_*)th rank as the distance threshold. Once the distance thresholds are obtained, we populate the matrix **V̅** = (*v̅_Im_*)_*N×M*_ by counting how many of the pooled distances from each structure *m* in the 2*M* distances are lower or the same as the corresponding distance threshold. Note that there are only three possible values of the matrix element: *v̅_lm_* = {0,1,2}. A value of 2 means that both homologues of TAD have to be located near the NE; a value of 1 means only 1 of the homologues has to be located near the NE; and a value of 0 means that neither homologue is forced to be located near the NE. The optimization step will then assign *v_im_* accordingly as either 0 or 1. When *v̅_lm_* = 1, the ambiguity as to whether (*v_im_* = 1, *v_i′m_* = 0) or (*v_im_* = 0, *v_i′m_* = 1) is solved on the fly, during the dynamic optimization of the genome structure, where 1 is favored for shorter distances to the NE.

### Optimization

As described elsewhere [22], we used step-wise optimization and the A/M iteration algorithm to generate the structure population. We first generated a population of structures satisfying all Hi-C constraints, then fine-tuned the model structures by gradually including the lamina-DamID constraints. For the Hi-C constraints, we included new contact probabilities in several stages during the optimization, at the lower thresholds **Θ** = {1, 0.7, 0.4. 0.2, 0.1, 0.07, 0.06}. One or more iterations were performed at every probability level. Contact probabilities less than 0.06 were not used at all. 26 A/M iterations were required to generate a structure population consistent with the Hi-C data. The lamina-DamID data were also included in several stages, at the probability levels **Θ** = {0.2, 0.1, 0.06}. Ten additional A/M iterations were performed to optimize the structure population with respect to the lamina-DamID data. The optimization was performed using a combination of simulated annealing molecular dynamics and conjugate gradient methods. The algorithm was implemented using the Integrated Modeling Platform (IMP) [57].

### Data collection and processing

Our processing methods for Hi-C, lamina-DamID and other epigenetics data are described in the Supplemental Material.

### Analysis of the structure population

Our statistical analysis of the structure population and details on all statistical tests are described in the Supplemental Material.

### Cell culture and immunofluorescence

Kc cells were maintained at 25°C as logarithmically growing cultures in Schneider’s medium (Sigma) + FBS (Gemini), and fixed and stained as previously described [46]. The antibodies used were anti-Fibrillarin (Cytoskeleton, Cat. #AFB01, 1:200) and anti-H3K9me2 (Upstate, Cat. # 07-442, 1:500).

### Larval fluorescence in situ hybridization (FISH)

Wild-type *w^1118^* flies were raised at 25°C. Brains were dissected from third instar larvae and squashed before fixation, as described in [58]. Fixation and FISH staining were carried out as described in [59], using the following probes: 5’-6-FAM-(AACAC)_7_ for chromosome 2 satellites, 5’-Cy3-TTTTCCAAATTTCGGTCATCAAATAATCAT for chromosome X satellites (359 bp), and 5’-Cy5-(AATAT)_6_ for chromosome 4 satellites. FISH probes were purchased from Integrated DNA Technologies, and designed as described in [58].

### Imaging and image analysis

All images were captured using a Deltavision fluorescence microscopy system equipped with a CoolsnapHQ2 camera, using 60x and 100x objectives and 10-12 Z stacks with Z-intervals of 0.2-0.4. Images were deconvolved with softWorx software (Applied Precision/GE Healthcare) using the conservative algorithm with five iterations. The distances between signals in 3D volume reconstructions of Kc cells or in individual Z stacks of larval tissues were calculated with softWorx. All distances were normalized to the nuclear diameter of their respective cells. Quantification of FISH signals in larval brains was limited to cells that displayed clear homologous pairing, defined as proximal or overlapping FISH signals for each probe.

### Additional files

Additional file 1: This .docx file contains the supplementary methods.

Additional file 2: This .docx file contains the following supplementary figures: S1-S10. Legends for these figures are presented under each figure.

Additional file 3: This .xls file constains three supplementary spreedsheets, each included as a separate tab: Suppl. Tables 1 (Summary of the Pearson’s correlation between contact probability from structure models and Hi-C experiment), Suppl. Table 2 (Summary of chromosomal territory index (TI) for individual arms and pairs of homologous arms.) and Suppl. Table 3 (The sphere size of structural units of model).

## Authors’ contributions

QJ, HT, KG and FA designed the 3D modeling methodology and parameterization with input from IC and XJZ. QJ and HT generated and analyzed the genome structure population, and QJ, HT and FA interpreted the results. XL and IC carried out FISH and immunofluorescence experiments and analyzed the results. QJ, HT, FA, IC and XJZ wrote the manuscript. All authors read and approved the manuscript.

## Acknowledgements

The work was supported by the Arnold and Mabel Beckman foundation (BYI program) (to F.A), NIH (U54DK107981-01 to F.A and X.J.Z. and NHLBI MAP-GEN U01HL108634 to X.J.Z), and NSF CAREER (1150287 to F.A.). F.A. is a Pew Scholar in Biomedical Sciences, supported by the Pew Charitable Trusts. This work was also supported by a Mallinckrodt Foundation Award and NIH R01GM117376 to I.C. We thank L. Delabaere for assistance with FISH experiments and for generating some of the FISH probes, and the Chiolo Lab for helpful discussions.

## Supplementary methods

### A. Hi-C data processing

#### A.1 Bin-level contact frequency

The sequencing data were downloaded from Gene Expression Omnibus under accession number GSE34453 [1]. We adopted the pipeline developed by Leonid Mirny lab [2] to process the Hi-C data. First, the two sides of each read were mapped to the Drosophila melanogaster genome (assembly dm3) independently using bowtie2 [3] with “very-sensitive” option. We truncated the reads to 20bp, and then remapped the non-mapped and multiple mapped reads by increasing truncation length with 5bp gradually. The truncating step significantly yields more double-sided mapped reads. 216,199,696 uniquely double-sided mapped reads are retained from the original 362,669,793 paired reads (with the mapping ratio at ~60%).

Then, the reads alignments for the artificial or non-informative contacts were filtered out, including self-ligation products, the products without ligation junction and PCR duplicates. After filtering process, 14,481,367 interactions are left for downstream analysis. Third, the valid double-sided alignments were used to construct the genome-wide contact matrix at 40k bin size, resulting in a 3012*3012 matrix. Before correcting the matrix for biases, we performed four types of bin-level filtering which will affect the normalization procedure. We removed the contacts between loci located within the same bin; removed bins with more than 50% are N’s in the reference genome; removed 1% of bins with low coverage; and truncated top 0.05% of inter-chromosomal counts which truncated values of the top 0.05% to be the same as highest value among the rest 99.95%. Finally, the iterative correction was performed on the filtered genome-wide contact map to get a normalized map, denoted as *F_K_* = (*f_ij_*)_*K*K*_.

#### A.2 Bin-level contact probability

The contact probability is defined as the probability for observing a given contact in the structure population. We define a threshold value *f^max^*, which defines the frequency at which a contact is formed in 100% of the structure population. We assume the contact frequencies that constitute any stable TAD can serve as reference where the interaction could exist in 100% of the cells. First, we register all contact frequencies inside TADs. Then we apply an R (CRAN statistical software) function *boxplot.stats* on this contact frequency set to get the extreme lower whisker of the boxplot and set it as the *f^max^*. Namely, in R

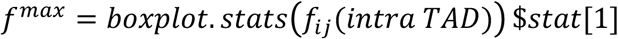

The contact probability between bin i and bin j is derived as

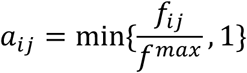

We applied the *f^max^* calculation method to the normalized frequency matrix to obtain the contact probability at 40 kb-binned matrix.

#### A.3 Domain-level contact probability

Based on the Hi-C contact frequency map, 1169 physical domains or topological associated domains (TADs) are detected by a quantitative probabilistic approach [1]. The chromatin in our population structure is represented at the level of TADs, therefore the contact probability at domain-level need to be obtained. The domain level contact probability is denoted as *A_N_* = (*a_IJ_*)_N*N_, where *a_IJ_* is the contact probability between domain I and domain J, and N is the total number of domains in the genome. *a_l_j* is derived from the corresponding contact probability at the bin level. If b(I) is the set of all bins in domain I, and b(J) is the set of all bins in domain J, then

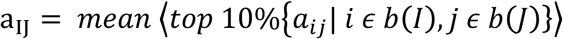

is the average value of the top 10% ranked contact probabilities in the set of all pairwise combinations between bins in *b*(I) and *b*(J).

### B. Lamina-DamID data processing

The genome-wide lamina-DamID binding signal is collected from [4]. The binding signal for each TAD (*L* = {*l_i_* = 1,2,…,*N*}) is calculated using BigWigSummary tool (from USC Genome Browser). The DNA content of the nuclear periphery was measured to be ~12% per nucleus for Kc167 cell line [5]. To reproduce the experimentally observed DNA content at the nuclear periphery, we relate the average lamina-DamID signal of all chromatin regions (measured in an ensemble of cells) to the average domain-NE localization probability ( *E* = {*e_I_*|*I* = 1,2,…, *N*}) in the structure population so that mean(*E*)=0.12. The lamina-DamID signal is transferred into a probability for each domain to be close to the NE as 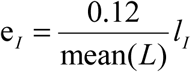.

### C. Analysis of the structure population

#### C.1 Reproducing lamina-DamID binding frequency

We define a chromatin domain-lamina contact, if the distance between the domain’s surface and the NE is less than 50nm. The NE-domain association probability is the fraction of structures in the population, in which the domain is in contact to the NE.

#### C.2 Chromosome territory index

To quantify how effectively one chromosome excludes other chromosomes from the volume it occupies in the 3D space, we adopted the quantity called chromosome territory. There is no universal definition of chromosome territory, but we follow the definition in a recent publication about the structure modeling of Drosophila polytene chromosomes [6]. Chromosome territory index (TI) is defined as the fraction of domains inside a convex hull that belongs to the chromosome used for its construction.

We first calculate the convex hull for a chromosome arm (with domain number *N_chr_*) using the function *delaunayn* in MATLAB (http://www.mathworks.com/help/matlab/ref/delaunayn.html). *T*=*delaunayn(X)* computes a set of simplices such that no data points of X are contained in any circumspheres of the simplices. The set of simplices forms the Delaunay triangulation. Then, the function *tsearchn* is used to search all the domains inside of the convex hull, and the number of detected domains is denoted as *N_hull_*. Finally, the TI is calculated as 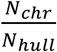.

The maximum *N_hull_* for each chromosome are the same, which is the total number of domains inside the nucleus (2238 euchromatin TADs in this study). The minimum *N_hull_* for each chromosome corresponds to the number of domains that belongs to themselves respectively. Under this definition, the maximum TI is 1, indicating that all domains inside the chromosome spanning volume are exclusively occupied by its own chromosome domains, and therefore shows a strong chromosome territory formation with only limited chromosome mixing. The theoretical minimum values for each chromosome arm are 0.096, 0.091, 0.095, 0.131, 0.008 and 0.0787, and for each pair of homologous arms are 0.192, 0.182, 0.189, 0.263, 0.016 and 0.157. The average TIs for chromosome arms in the structure population are 0.64 (2L), 0.65 (2R), 0.62 (3L), 0.62 (3R), 1.0 (4) and 0.67 (X). The average TI for individual arms is around 60%, suggesting the homolog pairs share territory almost equally. Indeed, the paired arms together possess high territorial index, i.e. 0.97, 0.98, 0.96, 0.98, 1.0 and 0.98 for arms 2L, 2R, 3L, 3R, 4, and X, respectively.

#### C.3 Residual polarized organization

The polarized (Rabl-like) organization shows that each chromosome occupies an elongated territory, with the centromere in one nuclear hemisphere and telomere in the opposite hemisphere. We investigated the position of each centromere and its corresponding telomere and obtain the number of polarized configuration chromosomes or arms (chr4 are excluded, therefore the number ranges from 0 to 10). To identify the presence of this organization, we measure the angle between each centromere, the nuclear center, and its corresponding telomere. If the cosine of the angle is positive, then centromere and telomere are in the same hemisphere. Otherwise, they occupy opposite hemispheres, forming polarized organization. If more than half of the chromosome arms (>=6) in one nucleus are in polarized organization, we consider this as a polarized nuclear structure.

#### C.4 Nuclear colocalizations of Hox gene clusters

The Antennapedia complex contains 5 genes located in 3 consecutive TAD domains, while the Bithorax complex contains 3 genes located in 2 TAD domains with one domain between them. Each control group contained two clusters, one cluster with 3 consecutive repressive domains and the other with 2 repressive domains separated by one domain. The 5 repressive domains were separated by the same linear distances as those in the Hox gene clusters. Because there are no available combinations of PcG TADs with the same genomic distances as the Hox gene clusters, the control data set involved the three types of repressive classes (Null, PcG and HP1 class). In total, we identified 30 combinations that meet the requirements of control groups. The average contact probability of each control group and hox gene clusters are shown in Fig. S5, top panel. All the contact probability of clusters are lower than 6%, which means the constraints are not imposed for those clusters in our models. If the closest surface-to-surface distance between two clusters in one group is less than 200 nm, we consider these clusters colocalized.

#### C.5 Pericentromeric heterochromatin cluster detection

We calculated all pairwise surface-to-surface distances (normalized by the sum of the radii of the domain spheres) among the 12 heterochromatin spheres, for each structure in the population, then obtained the average pairwise distances in the matrix. Hierarchical cluster analysis is performed on the average distance matrix by using *hcluster* function in R.

#### C.6 Homologous pairing

A domain is defined as paired if the surface-to-surface distance between two homologs is less than 200 nm. Then the pairing frequency is defined as how often a domain is paired among the structure population. The domains with frequency higher than Top 3^rd^ quantile are determined to be “tight”, otherwise, to be “loose”. We have 71 “active-tight” domains and 423 “active-loose” domains.

##### Pairing is not determined by the positon and the neighborhood crowdedness in the nucleus

The variation of the distance along homologous chromosomes and among nuclei raises the question of why certain regions attain higher level of homologous pairing than others in certain nuclei. First, we notice that the linear distance to centromere or to heterochromatin does not influence the extent of pairing of a domain (Fig. 6B), which exclude a possible influence of heterochromatin clustering on the extent of pairing. Next, we tested whether the 3D position of a domain in the nucleus influences pairing. We hypothesize that genomic regions near NE may have less space for movement, thus promoting homologous pairing. Indeed, the Pearson’s correlation between the contact frequency with NE (if the surface distance to NE is less than 50 nm, this domain is defined in contact with NE) and the homologous pairing frequency is 0.34 with p-value < 2.2e−16. Similarly, Pearson’s correlation between the average radial position and the homologous pairing frequency is 0.10 with p-value = 0.00049. Finally, we tested the hypothesis that the crowdedness of the neighborhood around a domain influences this domain’s pairing. The Neighborhood Crowdedness (NC) of a domain is defined as the number of other domains whose surface-to-surface distance to the domain is less than 200 nm. We calculate NC for each pair of homologous domains in individual models and compare the difference between paired and unpaired groups. The domains of paired groups have higher NC than the unpaired in 16.62% of models and lower in 6.84%, the rest of models (76.54%) show no significant difference. This data support the idea that the NC around domains does not significantly influence the pairing of homologous regions.

#### C.7 Epigenetic analyses

Chromatin domains were classified into four classes based on their epigenetic signatures: Active, Polycomb-Group (PcG), HP1/Centromere and Null [1]. Active domains comprise 42% of the domains with smaller domain size, and they are actively transcribed and characterized by high gene density. PcG domains are bound by PcG proteins and associated with the histone mark H3K27me3. HP1/Centromere domains are bound by the heterochromatin proteins HP1 and Su(var)3-9 and associated with H3K9me2. Null domains are not enriched for any of those marks.

We followed this 4-class annotation in our structure analysis. We also collected the data of histone modifications and binding of chromatin proteins in the study [4] from http://research.nki.nl/vansteensellab/Drosophila_53_chromatin_proteins.htm. Wig files were downloaded, and then transferred into bigwig format. BigwigSummary program (from USC Genome Browser) was used to extract the individual signals for requested regions (the defined 1169 TADs in this study). The signal was calculated as an average per domain, avoiding the bias of the genomic length.

#### C.8 Transcription analyses

Gene expression data (embryonic samples collected at 16-18h) were obtained from the modENCODE project [7]. 1169 physical domains covered 12947 genes with available expression data. The number of genes in each domain varies, ranging from 0 to 170, and the average number is 11. The average gene expression values are calculated for each domain. RNA polymerase II binding data for Kc167 cells were also from modENCODE project (accession no. GSE20806, link http://www.ncbi.nlm.nih.gov/geo/query/acc.cgi?acc=GSE20806). TBP (TATA-binding protein) is acomponent of the basal transcription machinery and the protein binding data are from [4]. BigwigSummary program was used to calculate the average signal for each defined domain.

#### C.9 DNA replication analyses

Data for ORC-binding regions and early activating replication origins for Kc167 cell line were downloaded from modENCODE (accession no. GSE20889 and GSE17285, respectively). BigwigSummary program was used to calculate the average signal for each defined domain.

#### C.10 Statistical test

The association test between two paired signals is done by *cor.test* function in R using Pearson’s product moment correlation coefficient. For example, the positive correlation between the frequency near NE derived from our population of structures and the lamina binding signal from lamina-DamID experiment provides a validation for our models; the negative correlation between the frequencies of homologous pairing derived from our population of structures and the Mrg15 protein binding signal from lamina-DamID experiment matches the unpairing function of Mrg15 protein and also validate our models.

The difference test between two sets is done by *wilcox.test* function in R, which performs a Wilcoxon rank sum test (equivalent to the Mann-Whitney test) when the population cannot be assumed to be normally distributed. For example, we found the ORC binding signals are much stronger in the active-loose than in the active-tight domains.

---

**Figure S1.**
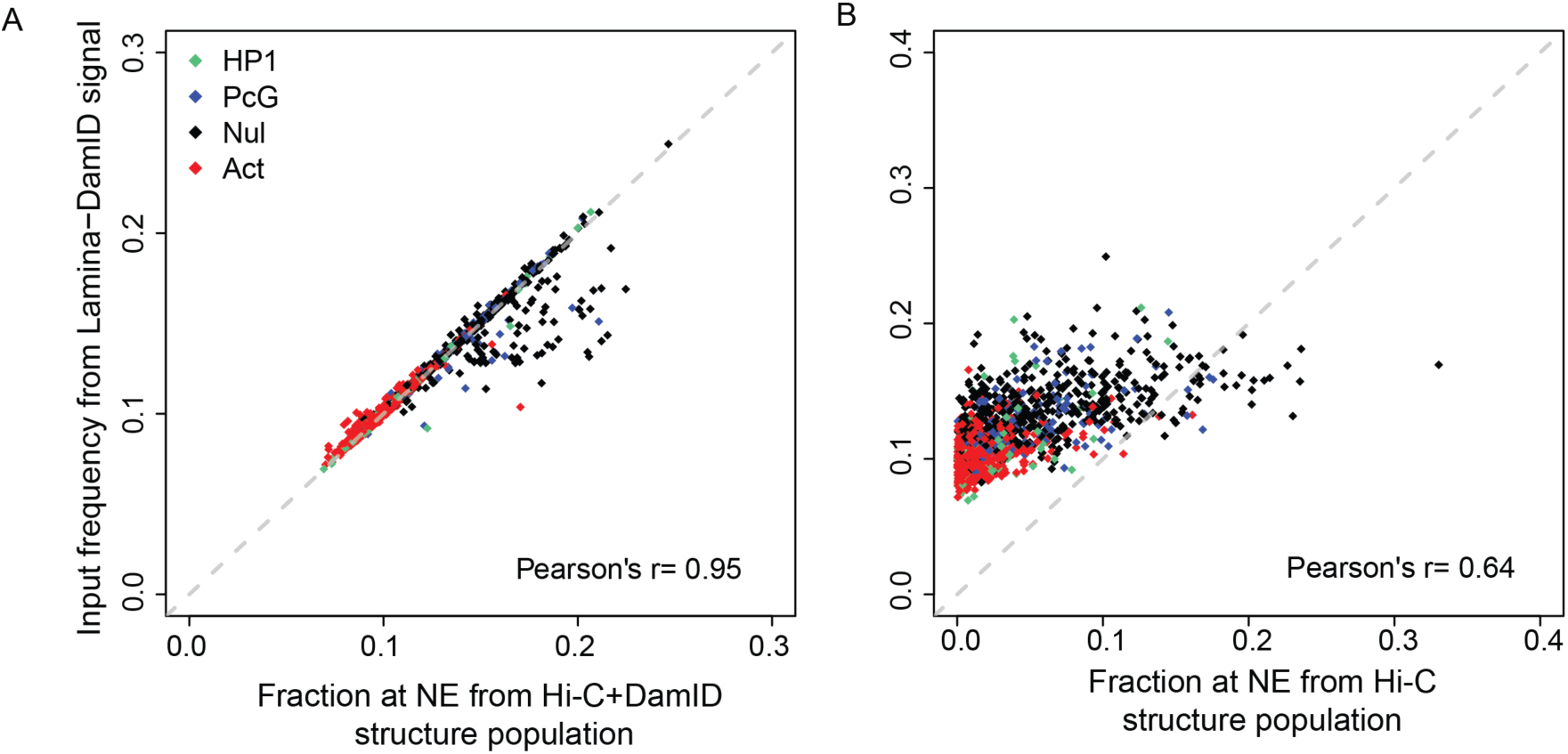
Agreement between the NE-association of euchromatin domains from lamina-DamID experiment and the models. (A) Fraction of domains at NE from population structure generated by data integration of Hi-C and lamina-DamID data well reproduces the input frequency derived from lamina-DamID data with Pearson’s correlation coefficient=0.95 and p-value< 2.2e−16. The points are colored according to the epigenetic classes. (B) Fraction of domains at NE from the control model with a structure population generated only from Hi-C data has a good correlation with the frequency derived from lamina-DamID data (Pearson’s correlation coefficient = 0.64 with p-value < 2.2e−16).

**Figure S2.**
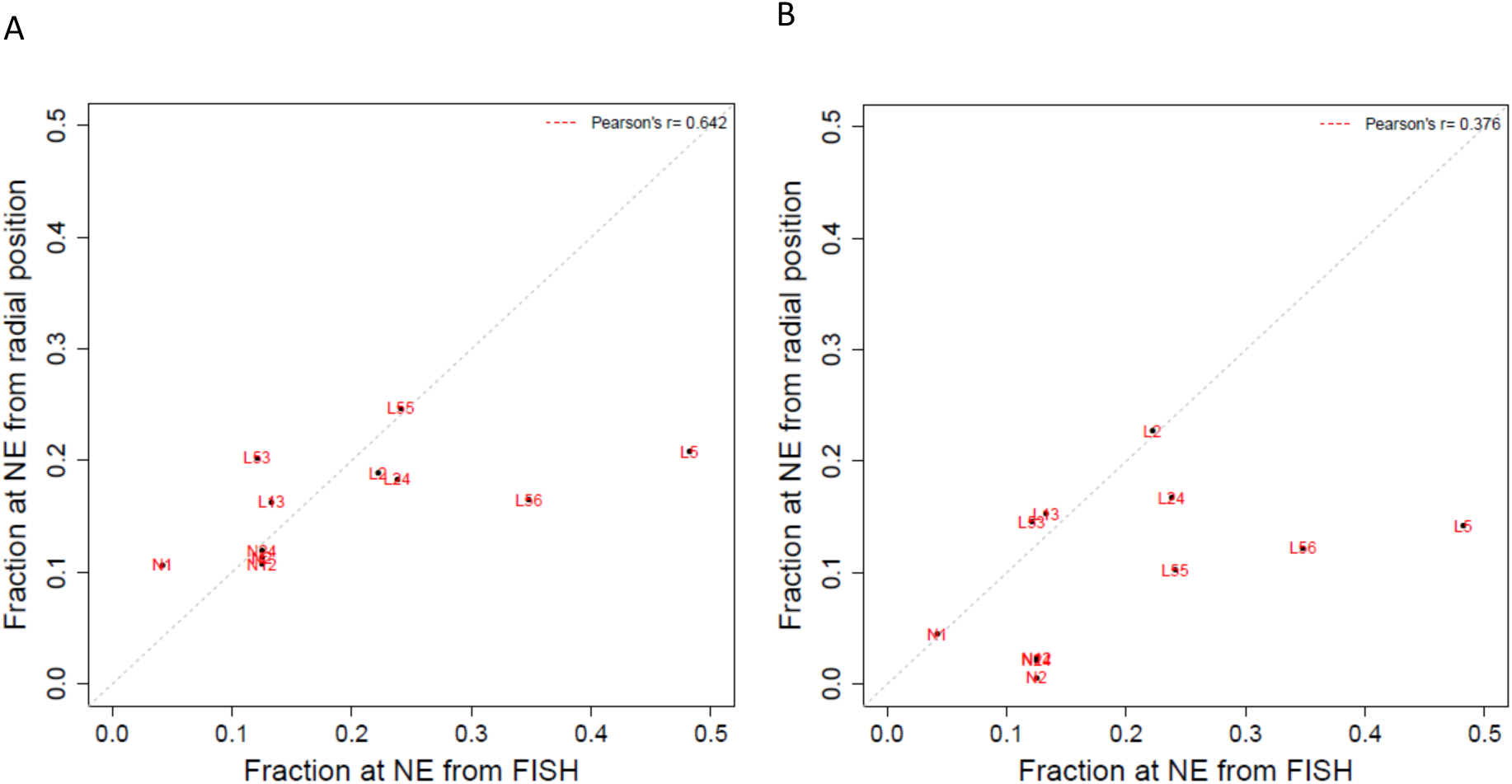
Agreement between the NE-association of individual loci from FISH experiment and the models. (A) Comparison of the NE association frequencies of individual loci from FISH experiment and from the model generated by data integration of Hi-C and lamina-DamID data. The NE association frequencies in the structure population agree well with FISH data for 11 loci (Spearman correlation coefficient=0.642 with p-value 0.03312). (B) Comparison of the NE association frequencies of individual loci from FISH experiment and the control model with a structure population generated only from Hi-C data. (Spearman correlation coefficient = 0.376 and p-value=0.2542).

**Figure S3.**
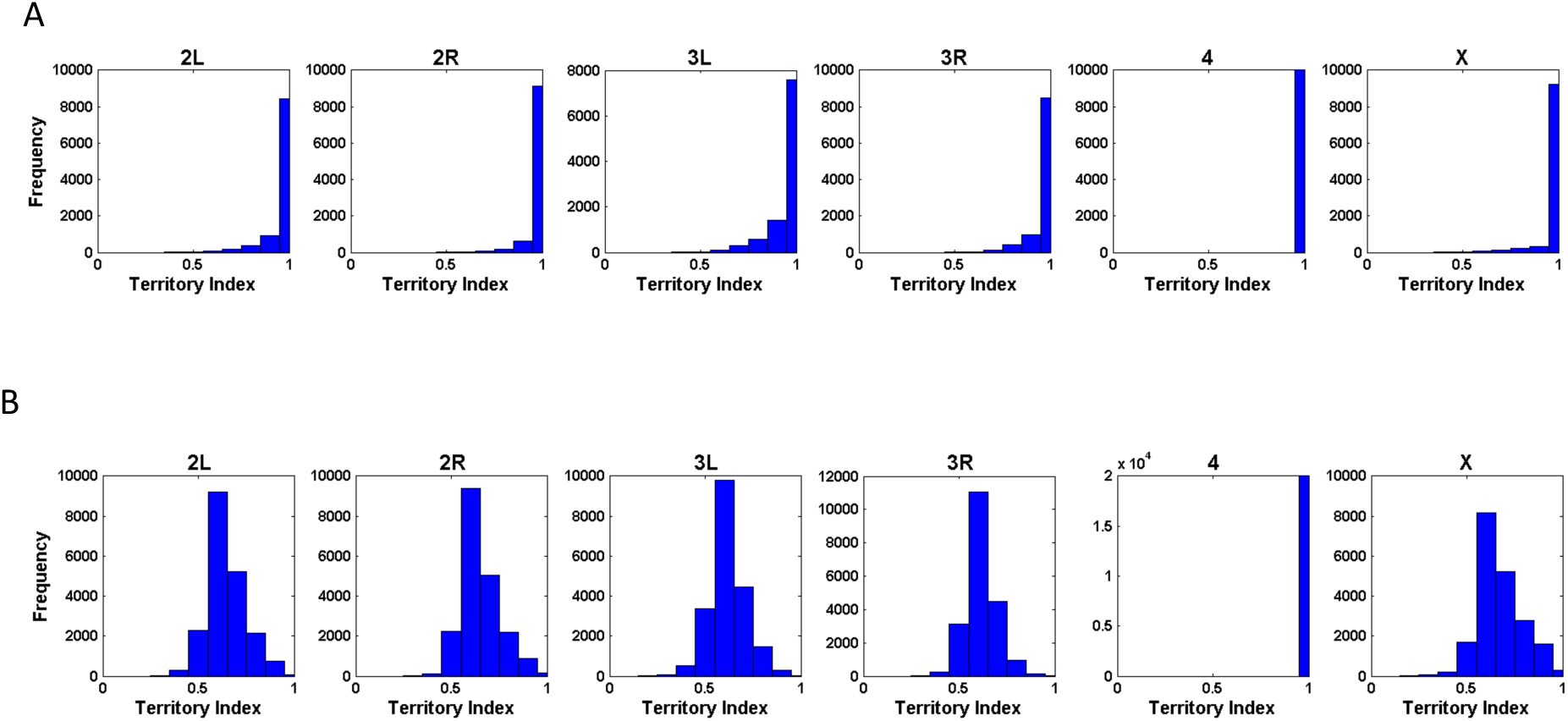
Territory index (TI). (A) TIs for the pairs of homologous chromosome arms. (B) TIs of each chromosome arm considering each homologues chromosome separately.

**Figure S4.**
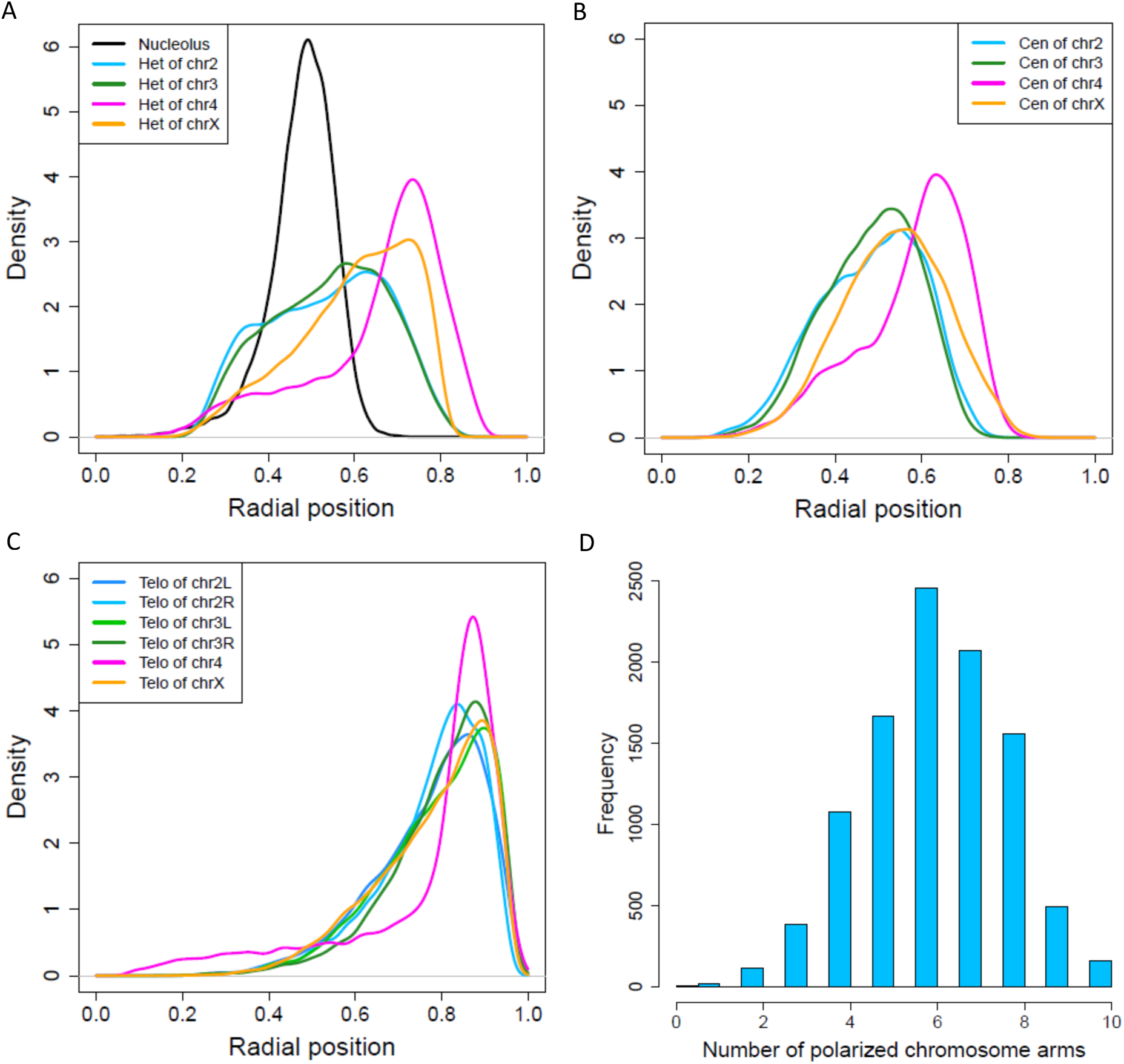
(A) Density plot of radial positions of the nucleolus and heterochromatin regions of different chromosomes. (B) Density plots of radial positions for centromeres. (C) Density plots of radial positions for peri-telomeric sequences. (D) Number of polarized chromosome arms (chr4 is excluded) among the population of structures.

**Figure S5.**
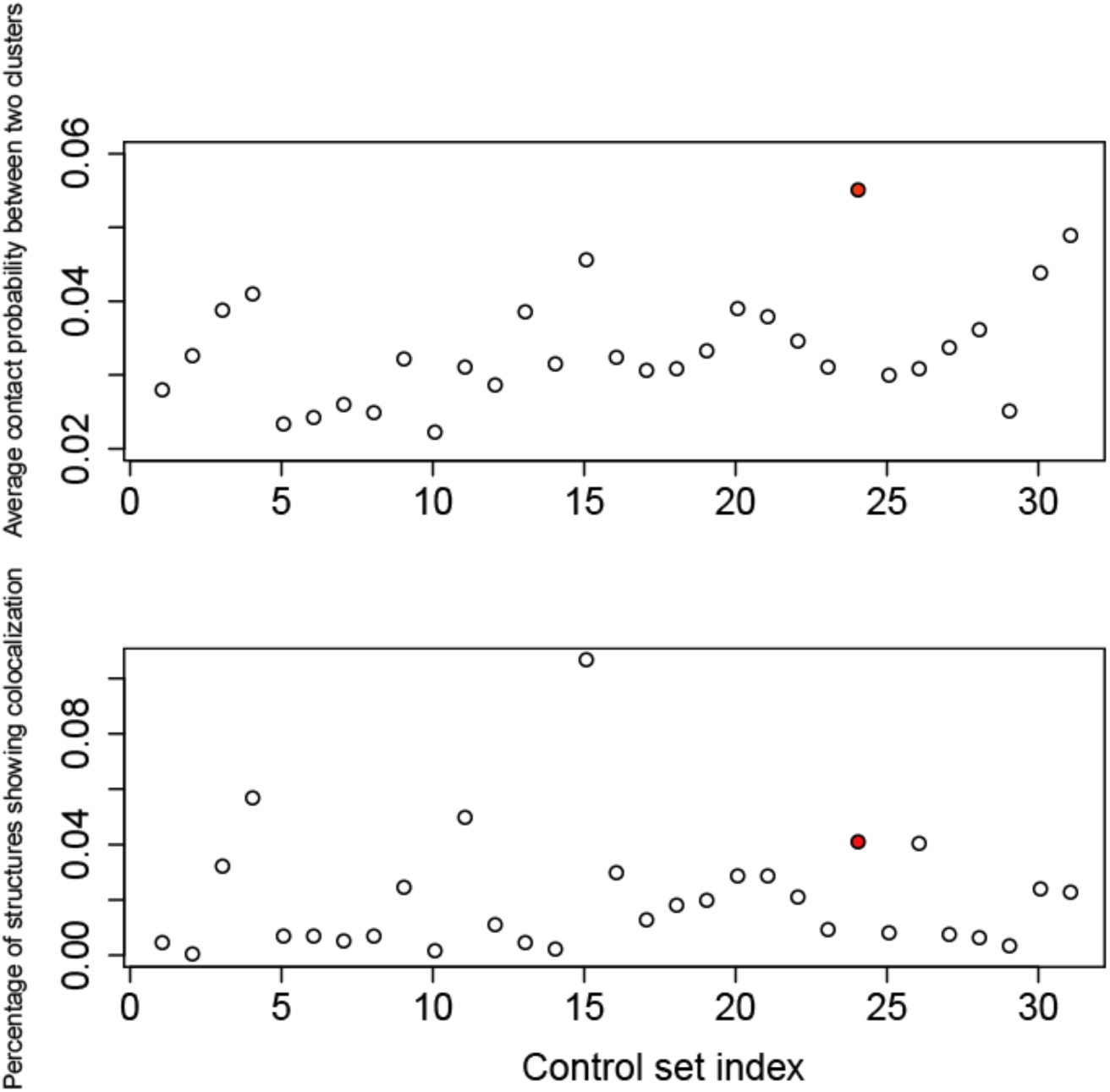
Hox gene clusters are prone to be co-localized comparing to control groups. (Top panel) Hi-C experiment shows the hox gene clusters have higher average contact probability than any other control clusters, and all the average contact probabilities within clusters (both hox gene and control) are smaller than 6%. The point for Hox gene clusters is highlighted by red color. (Bottom panel) The hox gene clusters show contacts in higher percentage of structures in the population compared to all other control clusters except three.

**Figure S6.**
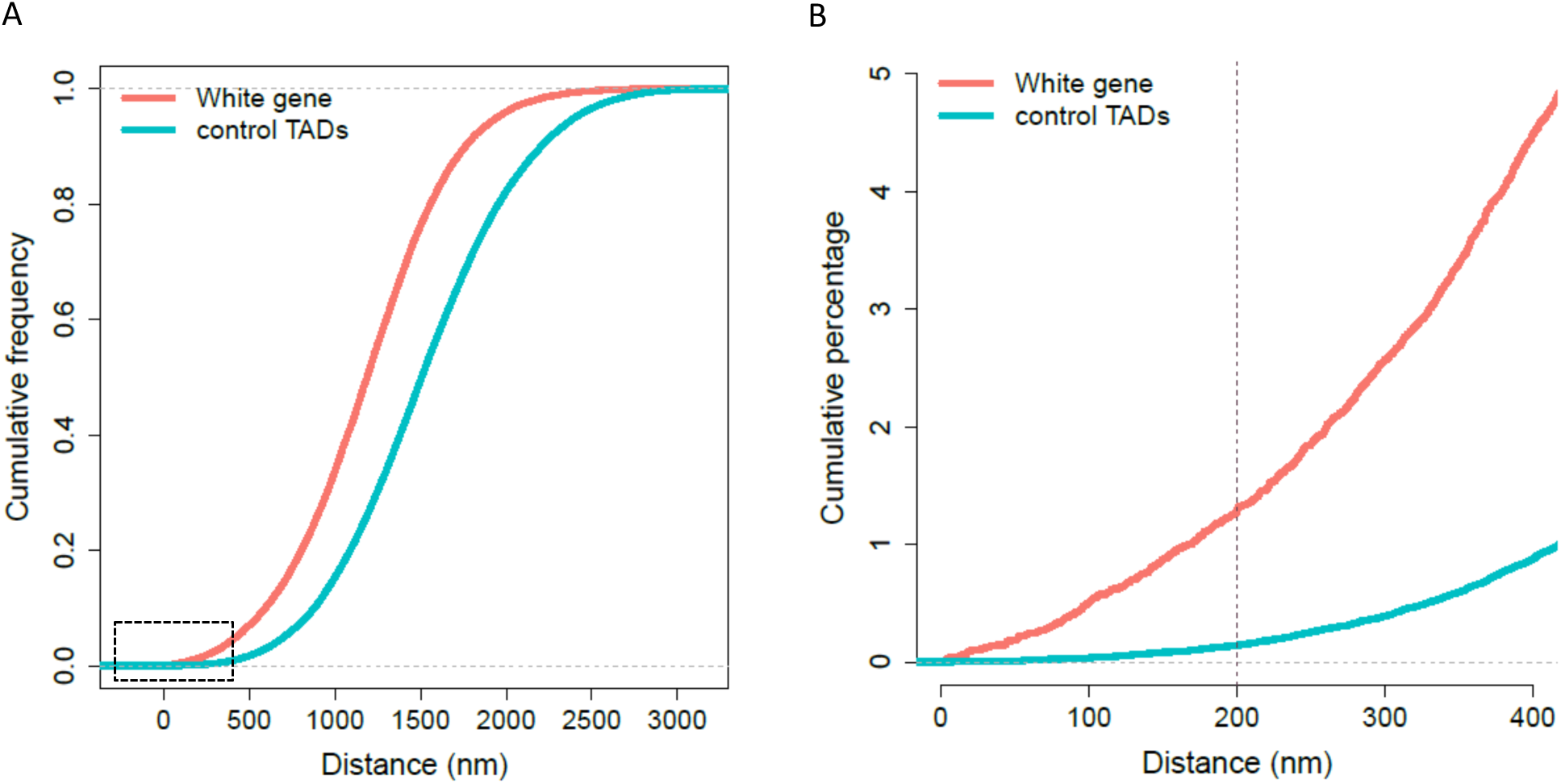
The *white* gene is prone to localize near to the pericentric heterochromatin. (A) Cumulative frequency plots for the distance of the *white* gene to its heterochromatin and of the control TADs to their corresponding heterochromatins. (B) Zoom the plot into the small distance, the *white* gene is 9-fold more frequently located proximal to pericentric heterochromatin (using 200 nm as a threshold), relative to control TADs.

**Figure S7.**
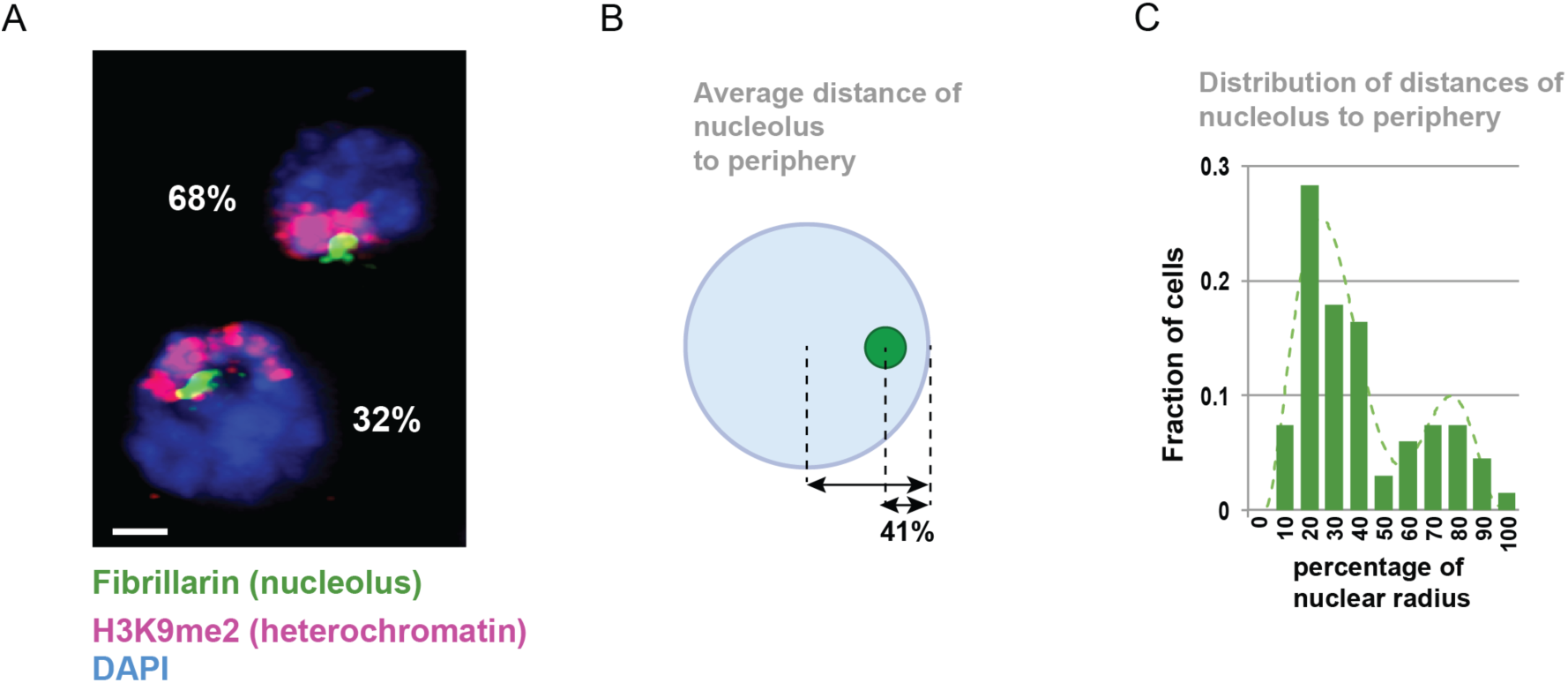
Nucleolus and heterochromatin positions in *Drosophila* Kc cells. (A) Immunofluorescence analysis with anti-Fibrillarin and anti-H3K9me2 antibodies, and DAPI staining for DNA (nuclear staining) shows the position and organization of the heterochromatin domain and the nucleolus in *Drosophila* Kc cells. The image shows a max intensity projection of two representative nuclei. Percentages indicate the population of cells in each configuration, *i.e.* with the nucleolus proximal to the nuclear periphery or more internal. N = 113 cells. Scale bar = 1 μm. (B) Quantification of the distance between the center of the nucleolus and the nuclear periphery shows the average position of the nucleolus relative to the center of the nucleus and the distribution of these distances in the cell population. N = 63 cells. (C) The distribution of the distance between the center of the nucleolus and the nuclear periphery show a bimodal distribution.

**Figure S8.**
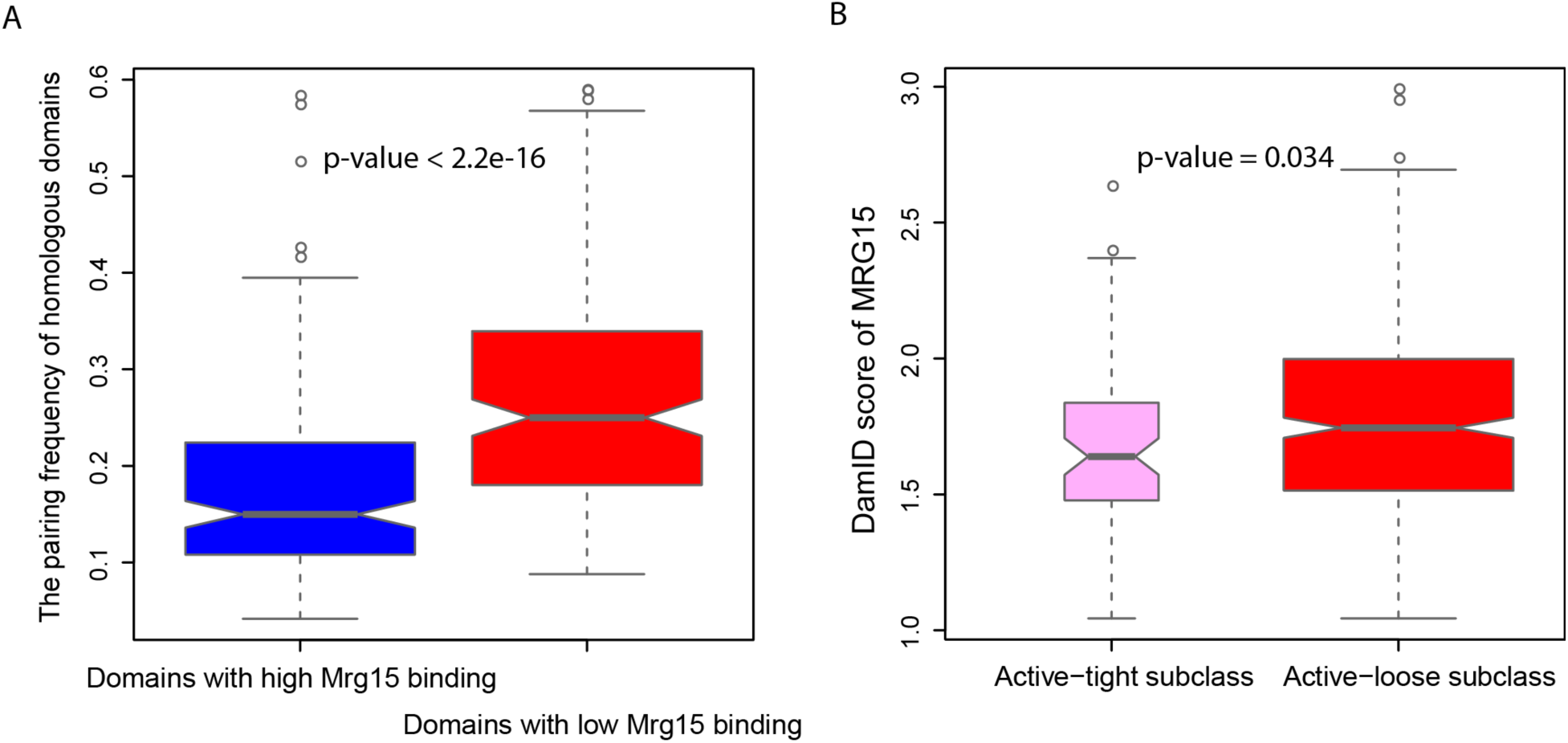
Inverse correlation between Mrg15 binding and homolog pairing frequency (A) Boxplot for the pairing frequency of domains with high and low Mrg15 binding. All domains are divided into 3 subsets based on their Mrg15 binding score. 293 domains are in the subset with high Mrg15 (top 25% binding scores); 293 domains are in the subset with low Mrg15 (bottom 25% binding score). The pairing frequencies for domains enriched with Mrg15 are significantly less than those for domains with low Mrg15 score (one-tailed Mann-Whitney U test, p-value < 2.2e−16). (B) Boxplot of Mrg15 score for the active-tight and active-loose subclasses. Active domains are divided into active-tight and active-loose based on their pairing frequencies (**Suppl. Methods C.6**). Active-tight domains, which have high pairing frequencies in our models, are significantly less enriched with Mrg15 comparing to active-loose ones (one-tailed MannWhitney U test, p-value = 0.03436).

**Figure S9.**
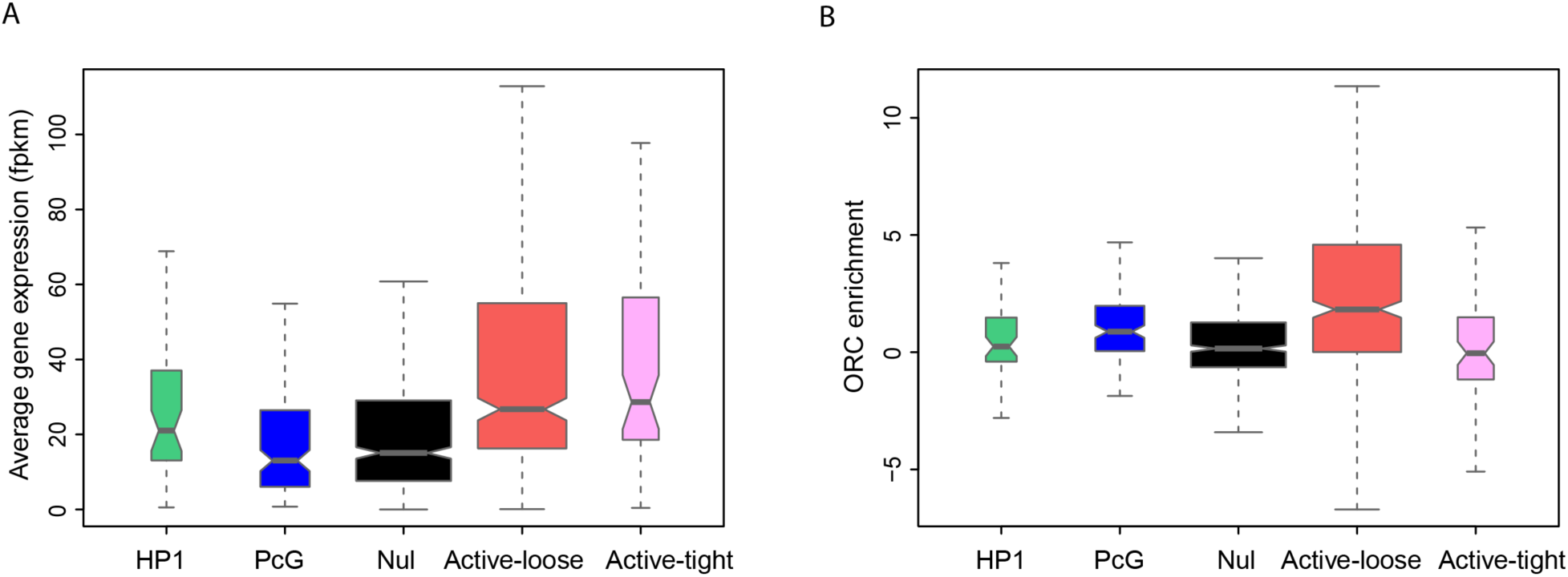
Transcriptional activity and DNA replication for all five classes (A) Domains in both two active subclasses have higher gene expression levels compared to those in the three repressive classes. (B) Domains in the active-loose subclass are more enriched with the replication complex ORC compared to the domains in the three repressive classes and the active-tight subclass. p-values: 1.48e-4, 7.59e-4, <2.2e−16 and 2.78e-5 respectively for HP1, PcG, Null and Active-tight.

**Figure S10.**
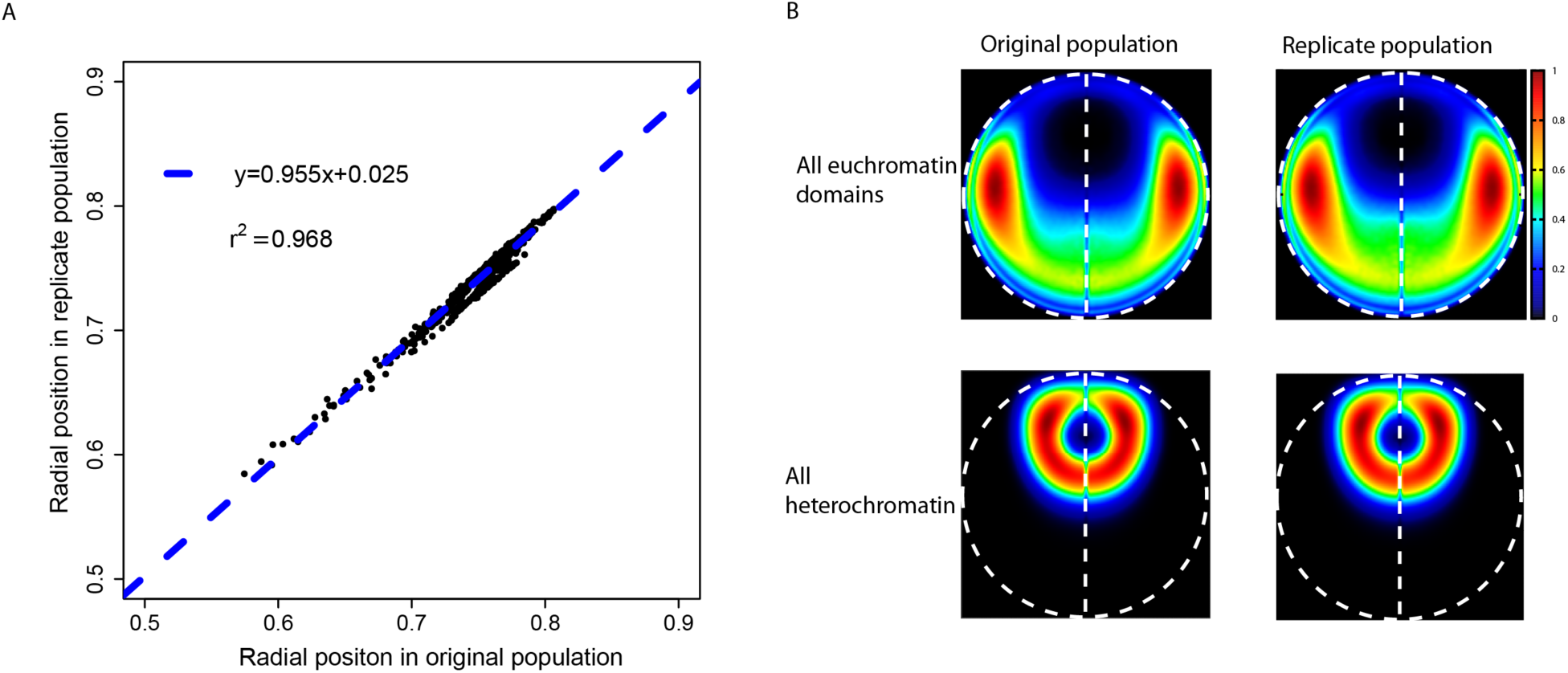
Reproducibility of the modeling method applied to the Drosophila genome (A) Agreement of the average radial positions between two populations of structures. The Pearson’ s correlation between them is 0.984, with p-value < 2.2e−16. (B) (Top panel) LPD plots of all euchromatin domains for the original population and the replicate population respectively. (Bottom panel) LPD plots of all heterochromatins for two populations of structures show highly consistent results.

